# Automatic binding of basic sensory features requires consciousness

**DOI:** 10.64898/2025.12.10.693567

**Authors:** Zhili Han, Hao Zhu, Qian Chu, Yuchunzi Wu, Xu Chen, Yuanqing Wang, Sixian Li, Xiangbin Teng, Patrick C. M. Wong, Chen Yao, Xing Tian

## Abstract

Conscious awareness requires establishing coherent percepts. Yet, whether consciousness is necessary for initiating the integration of basic sensory features remains unclear. Competing theories implicate distinct functional regimes of consciousness in the process of feature binding and creating conscious percepts. We used a novel multi-feature oddball paradigm with intracranial stereo-electroencephalography (sEEG) recordings in awake and anesthetized states to investigate the functional boundary of consciousness. In the awake state, the auditory attributes of loudness and tone, as well as the binding of the two features, were automatically encoded without attention to the stimuli in a functionally localized sensory cortical network. In the anesthetized state, the cortical registration of single attributes was preserved, whereas the binding was abolished. Moreover, anesthesia mostly influenced later cortical processes after stimuli offset. These results reveal the borderline of exertion of consciousness between the encoding and manipulation of basic sensory features in local cortical circuits – the functional boundary of consciousness constrains the feedforward binding and recurrent process directly at local rather than global level computations.

## INTRODUCTION

Our conscious experience requires the establishment of coherent percepts by integrating sensory features of the same object^1,2^. The encoding of basic sensory features can be pre-attentive and automatic without awareness^3,4^. Whereas, the integration of features requires task-driven top-down attention^5,6^, suggesting a possible functional boundary of consciousness between encoding and manipulation of sensory features in creating conscious percepts.

Competing theories implicate such distinct borders between functions in which necessitate consciousness (functional boundaries of consciousness). The critical stage that mediates the emergence of consciousness has been postulated as computations at the global or local level^7^. For one group of theories, such as the integrated information theory^8,9^ (IIT) and the global workspace theory^10,11^ (GWT), information needs to propagate and compute in global associate regions so that consciousness can be ignited. Whereas for another group of theories, such as higher-order theories^12,13^ (HOT) and re-entry theories^14^, the key computations at local sensory regions are sufficient to give rise to consciousness. Different groups of theories would predict the operational threshold of consciousness between encoding and binding of sensory features at either the global or the local level. The functional boundary of consciousness is an informative and ubiquitous factor in all theories, providing an inclusive and unbiased investigation by taking all types of theories into account^15^.

Evaluating the functional boundary of consciousness is complicated by the nature of the methods. Directly observing the neural correlates of consciousness with introspection and self-report can provide phenomenal characters of consciousness^16,17^. However, it is hard to assess the emergent process and impossible to distinguish between the correlates to the consciousness itself or to the reporting process^7,18–20^. By contrast, general anesthesia offers a ‘temporary lesion’ of consciousness. It suppresses consciousness while leaving reporting mechanisms irrelevant, enabling contrapositive tests of emergence and boundary conditions. Convergent evidence shows state-transition markers in EEG during propofol-induced loss to recovery, disruptions of fronto-parietal integration, and systematic reductions in brain-signal complexity. Together, these provide quantitative, report-free indices of the functional boundaries of consciousness^21–25^. Recent syntheses integrate these findings with leading theories, clarifying how anesthesia specifically disrupts conscious processing^26^.

The Oddball procedure is an optimal experimental paradigm for quantifying the functional boundary of consciousness. A stimulus that is deviant from the regularity of the preceding sequence can induce a prominent neural marker of mismatch negativity (MMN), indexing the establishment of a novel neural representation^27–29^. The MMN has been observed in the absence of attention and during sleep^30–32^, suggesting an automatic process without awareness. Moreover, MMN to the deviant of a single sensory feature has been observed even under anesthesia^33^, providing the minimal neural computation that does not require consciousness and hence the potential lowest cutoff of the functional boundary^34^. Recently, a multi-feature oddball paradigm was developed and was used to quantify the automatic feature binding process^35^, which is an ideal paradigm to probe the functional boundary of consciousness -- whether the conscious state is a prerequisite for automatic feature binding.

To test whether automatic feature binding necessitates the conscious state, we probed the temporal dynamics using the multi-feature oddball paradigm in a within-subject design directly contrasting the two conscious states of wakefulness and general anesthesia^33,36,37^. Neural responses were recorded using intracranial stereo-EEG in functionally localized sensory regions for basic auditory attributes pre-defined using fMRI. The integrative process is widely considered a key substrate of conscious awareness^14,38,39^, and is mediated by both feedforward and recurrent dynamics between neural populations^35,40–42^ in a temporally synchronized manner^43,44^. If the functional constraints of consciousness lie between the encoding and binding of basic sensory features in the local sensory network, MMN responses to single-feature deviants would survive anesthesia, whereas responses to multi-feature deviants would be selectively abolished in dedicated integrative regions of auditory cortices. The existence of a borderline would indicate the functional boundary of consciousness constraining feedforward binding and recurrent processes in a local sensory cortical network (Fig. 1A).

**Figure 1.**
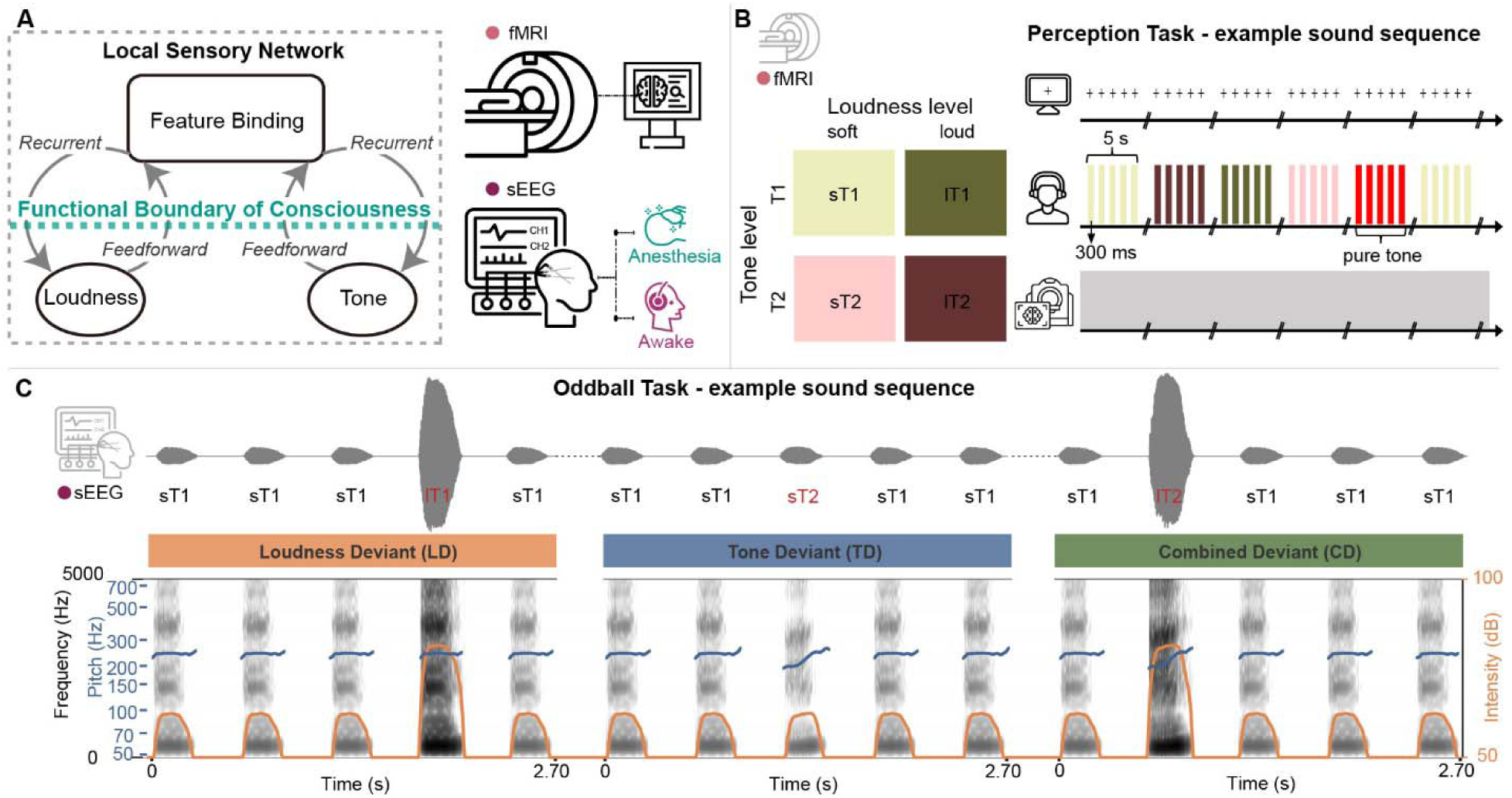
Theoretical schematic and experimental procedures in multimodal investigation on how the conscious state constrains auditory feature binding. A) Theoretical schematic of the functional boundary of consciousness. The functional borderline of consciousness is hypothesized as between the encoding of auditory attributes and the binding of sensory features, by constraining both feedforward and recurrent processes in a local sensory network. Functional MRI (fMRI) scanning was used to delineate the cortical distributions of processing basic auditory features. Intracranial stereo-electroencephalography (sEEG) was recorded in both conscious (awake) and unconscious (general anesthesia) states to probe the functional boundary of consciousness in the auditory cortical hierarchy. B) Experimental design of the fMRI study for localizing the functional sensory regions that represent auditory attributes. Left: auditory stimuli were created from a full combination of two auditory attributes: loudness (soft vs. loud) and Mandarin tone (first tone vs. second tone, T1/T2), forming four distinct categories. Right: In a mini-block design, each stimulus category was presented five times (stimulus duration of 300ms, stimulus-onset asynchrony, SOA of 1s). Mini-blocks were separated by a 10 s silent interval. Randomly inserted pure-tone probes (red bars) served as catch trials for verifying participants’ attention in the perception task. C) Experimental design of the sEEG study for probing automatic feature binding. A multi-feature oddball paradigm was used during sEEG recordings. Patients were presented with auditory sequences containing three types of deviants (labeled in red) from a more frequently occurring sound (standard). The example showed with sT1 sound as the standard, and the changed features in deviants were depicted the features of changes below. Stimuli were 300ms in duration with a fixed 300ms inter-stimulus interval. The paradigm was administered during both the intraoperative general anesthesia state and the postoperative awake state. (See Table S1 for details.)

## RESULTS

### Multivariate fMRI decoding reveals distinct cortical maps for loudness and tone

To reveal the distributions of neural representations of basic auditory attributes as anatomical bases for sEEG investigation of feature binding, fMRI data were recorded when participants listened to two Mandarin tones with two levels of loudness (Fig. 1B). The behavioral performance of detecting a pure tone target was at ceiling, indicating that participants were engaged in the perception task. Multivariate decoding was applied to the fMRI data. The main effects of classification were significant for loudness (*t*(19) = 26.784, *p* < 0.001, Cohen’s *d* = 5.99) and tone (*t*(19) = 29.522, *p* < 0.001, Cohen’s *d* = 6.60), demonstrating the sensitivity for revealing robust and feature-selective representations for both loudness and tone.

For the attribute of loudness, the most reliable decoding was observed in bilateral Heschl’s gyrus (HG, core auditory cortex) and extended into the left planum temporale and superior temporal sulcus (Fig. 2A and Table S3), consistent with rapid encoding of envelope-based salience^45,46^. For the attribute of tone, decoding results preferentially revealed in more lateral and anterior areas, including the planum polare and anterior superior temporal gyrus (aSTG), as well as the right angular gyrus (Fig. 2B and Table S4), consistent with phonological–semantic integration of lexical tones^47–49^.

**Figure 2.**
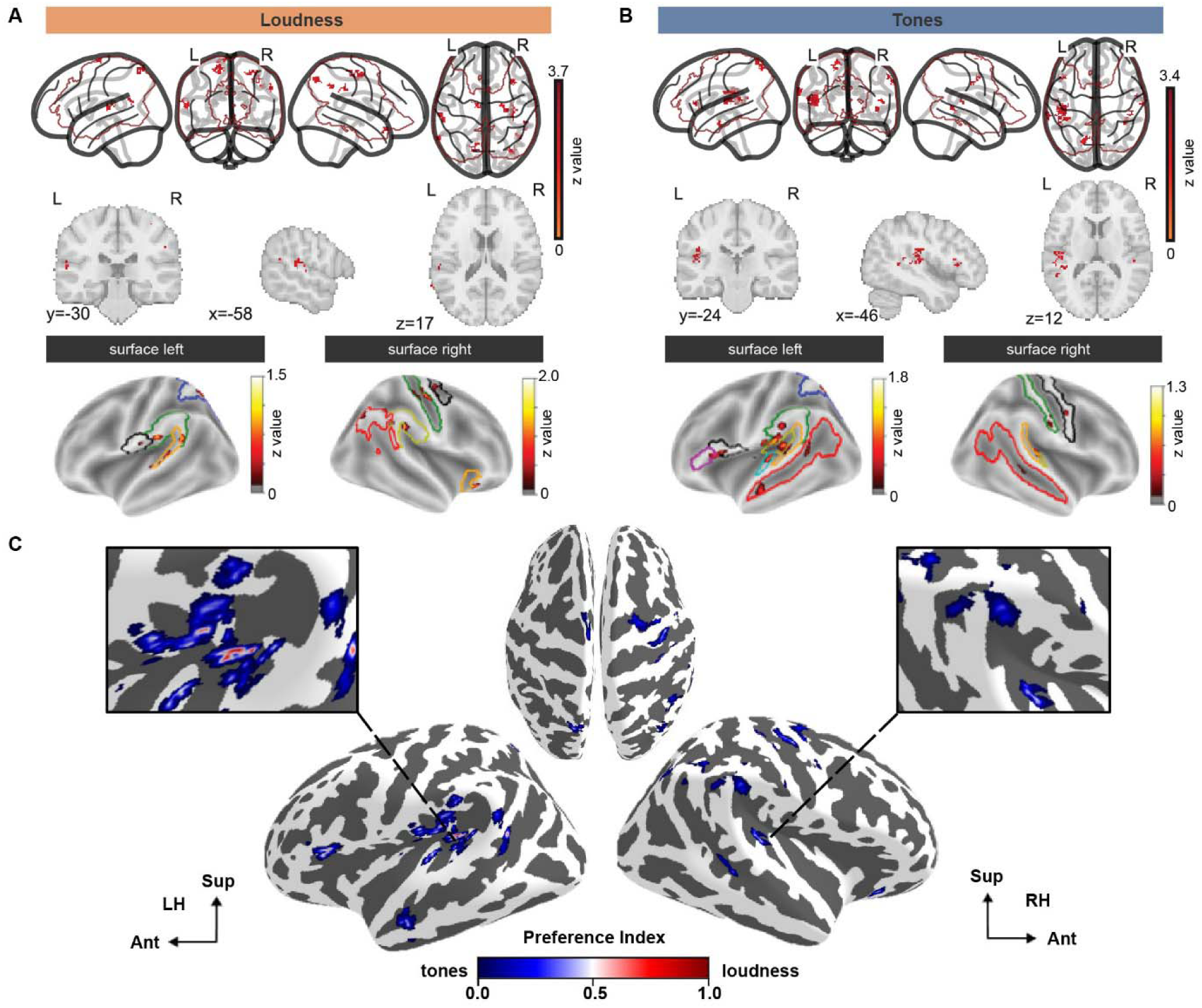
Results of fMRI decoding reveal distributed cortical regions for distinct and overlapped representations of loudness and tone. Multivariate decoding of BOLD responses revealed distributed cortical representations for A) tone and B) loudness. Decoding accuracies were derived from searchlight-based linear discriminant analysis (LDA) across auditory-related cortical parcels. Robust decoding of loudness was centered on the core auditory cortex, including bilateral HG and extending into the left planum temporale. Lexical tone decoding was robust in lateral and anterior regions, including the planum polare and anterior superior temporal gyrus (STG). (See Table S2 for details.) C) Results of preference analysis. A preference index was derived based on the weighting of loudness and tone decoding and revealed a center-surrounding spatial gradient (medial loudness-preference to lateral and anterior tone-preference) in posterior HG and middle STG regions.

We carried out a preference analysis to further reveal the cortical gradient of feature representation (Fig. 2C). A preference index was calculated by the relative weighting of decoding scores for loudness and tone. The distribution of the index illustrated that loudness-selective regions were concentrated medially and centrally, particularly in the core HG and adjacent planum temporale. Whereas the tone-selective regions surrounded the loudness-selective zones laterally and anteriorly, encompassing the belt regions of HG, planum polare, and aSTG. Beyond the auditory cortex, loudness selectivity extended into the inferior frontal gyrus and supramarginal gyrus, aligning with dorsal and ventral pathways for speech processing^47,50^, whereas tone selectivity uniquely recruited the right angular gyrus and ventral sensorimotor cortex, consistent with prosodic and articulatory integration^49,51^. Crucially, overlapping selectivity was observed in left lateral HG and posterior planum temporale. The joint representation regions are potential candidates as integration hubs where individual feature channels converge. The fMRI decoding results demonstrate that the attributes of tone and loudness are represented in spatially adjacent and partially overlapped cortical areas, providing target anatomical substrates for electrode selection in the subsequent sEEG investigation of feature binding.

### Double dissociation of repetition effects in early and late process stages between conscious states

To investigate how conscious states constrain automatic feature binding, we conducted a multi-feature oddball experiment during both awake and general anesthesia in the same participants. Sequences of sounds were presented, in which an identical sound was played several times in a row (standard) before a different sound (deviant) occurred (Fig. 1C). To establish validity, we first examined the high-frequency activity (HFA, 70–150 Hz) of the auditory response elicited by the standard stimuli, collapsing across different types of sound. Auditory stimuli evoked robust high-gamma responses in all participants (Fig. 3A), localized predominantly in bilateral auditory cortices, most consistently in left HG and adjacent superior temporal gyrus (STG) (Fig. 3B). This spatial distribution matched the fMRI decoding results (Fig. 2C), providing convergent evidence that sEEG captured the same early processing of basic auditory attributes. The following analyses concentrated on the functionally defined auditory contacts.

**Figure 3.**
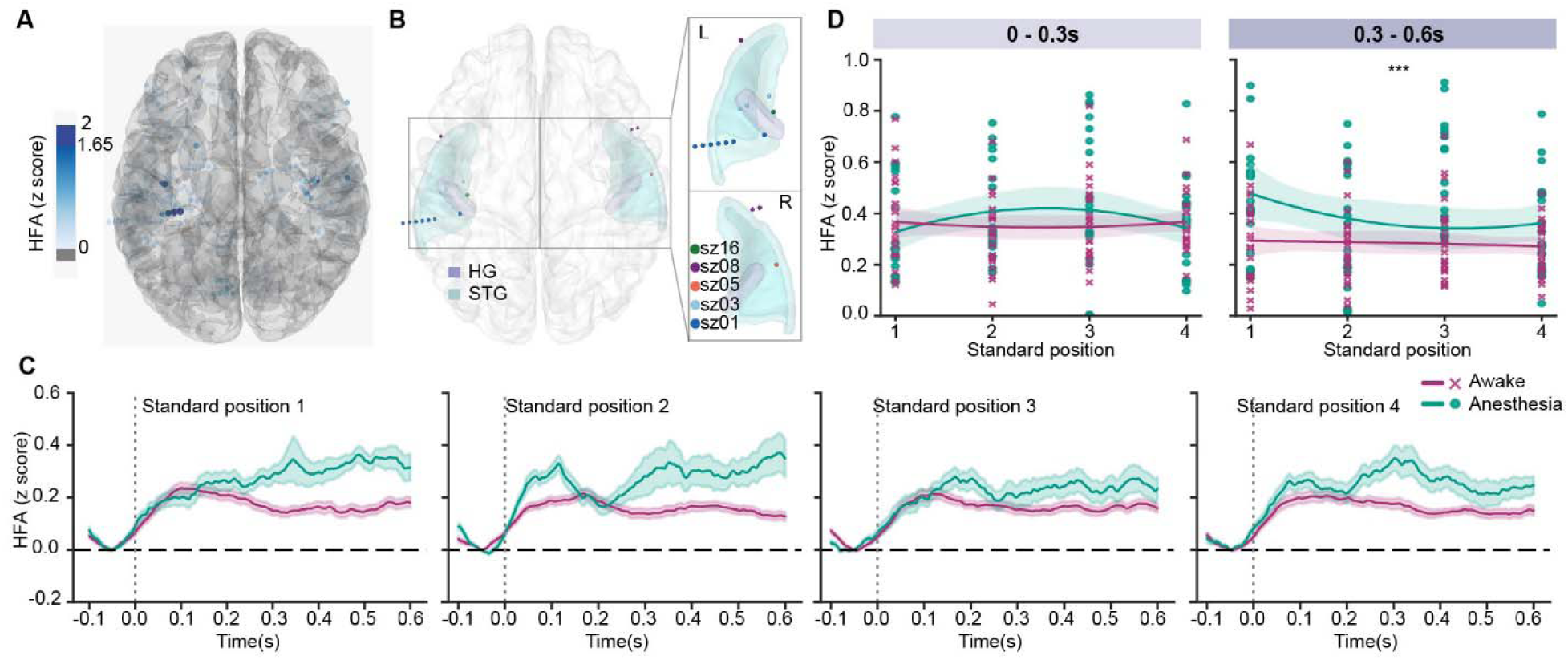
Loss of consciousness disrupts the repetition effects in later but not early sensory processing. A) Electrode contacts of sEEG that show responses to auditory stimuli in the awake state. The response strength is color-coded, with deep blue denoting significant contacts (*z* > 1.65, *p* < 0.05). The significant contacts were mostly in bilateral auditory cortices, with the strongest responses localized to the left Heschl’s gyrus (HG) and superior temporal gyrus (STG). B) Functional and anatomical jointly defined auditory contacts used in analysis. Contacts were selected based on significant HFA responses to auditory stimuli in the awake state in (A) and further restricted to anatomically defined auditory cortex using Desikan-Killiany and Destrieux atlases. A total of 15 contacts from 5 participants were included. C) Neural responses to the four consecutive standards (S1-S4) in awake (red) and anesthetized (green) states. The HFA responses to standard stimuli were averaged across trials and auditory contacts. The sound onset was at 0 ms and offset at 300 ms, followed by a 300 ms blank period. In the awake state, a peak around 0.1 s was observed in all four positions, followed by responses at a lower plateau. The early peak resembles a canonical auditory response; the peak and responses in the following time period remain at a minimal level, indicating the repetition suppression effects caused by repeated presentations of the standard stimuli. In the anesthesized state, the early auditory peak was comparable to that in the awake state. However, the responses elevated and diverged from those in the awake state, suggesting that anesthesia influences later processing in the sensory region. Shaded areas represent SEM. D) Statistical quantification of early and late processes in two conscious states. The HFA responses were averaged in the early (0 to 0.3 s) and late (0.3 to 0.6 s) time windows, and plotted as a function of standard position (S1–S4). No differences were observed between awake and anesthesized states in the early window (left). Whereas in the late window (right), responses in the awake state were significantly lower than those in the anesthesized state (p = 0.001), suggesting that the loss of consciousness influences local recurrent process in sensory regions in a later processing time. (*p =< 0.05, **p =< 0.01, *** p =< 0.001).

Next, we examined the repetition effects of the standard stimuli in the awake state, a neural hallmark of establishing regularity. In the 15 functionally defined auditory contacts localized in the STG and HG (Fig. 3B), HFA elicited by four sequential standard positions (S1-S4) rapidly attenuated and stabilized at a low level of activation throughout the sound presentation period (0-0.3s) and after the sound offset (0.3-0.6s) (Fig. 3C, red lines). These results are consistent with common observations of repetition suppression^52–55^, indicating successful formation and maintenance of regularity in auditory sequence.

We next tested the effects of general anesthesia on the processing of basic auditory attributes. First, we objectively validated the manipulation of the conscious state by quantifying the aperiodic 1/f spectral slope from the power spectral density estimated in all electrode contacts^56^. Spectral slopes were significantly steeper during anesthesia than wakefulness (*t*(12)= –3.13, *p* = 0.009) (Fig. S3), consistent with reduced high-frequency activity and cortical arousal in the loss of consciousness during anesthesia^56^. Moreover, despite weaker auditory responses during anesthesia compared to awake (Fig. S4; *p* = 0.022, cluster-based permutation tests, significant cluster window 62–142ms), the responses in minimal conscious state still have a significant peak with a relatively similar dynamic profile as those during the awake state, suggesting that basic auditory features can still be registered even without consciousness.

When evaluating how repetition effects of standard stimuli were influenced by anesthesia, a dissociation between early and late neural processing was observed (Fig. 3C). Specifically, during the early window (0∼300ms) when sound was presented, the magnitudes of HFA responses were similar between the awake and anesthetized states. However, in the later window (300∼600ms) when the sound was offset, the responses in the anesthetized state drifted up in all four positions of standard stimuli. A linear mixed-effects model (Fig. 3D) revealed no significant model effect in the early window (R^2^ = 0.002, F(3,188) = 0.127, p = 0.944), indicating no difference in the early sensory processing between the two consciousness states. In contrast, a state-dependent effect emerged in the late window (R^2^ = 0.081, F(3,188) = 5.547, p = 0.001), where a significant difference between awake and anesthesia states was observed (t(188) = 5.746, p < 0.001), suggesting that general anesthesia disrupted the repetition effects in the later sensory processing. Specifically, under anesthesia, the HFA responses decreased as the repetition of standards increased and plateaued toward the end (*β* = −0.038, *p* = 0.048). The double dissociation in early and late processing stages between two conscious states suggests that the repetition effects in the feedforward process in the early window were largely preserved in the unconscious state, whereas the recurrent process in the late window was absent because of anesthesia. These results are consistent with the local recurrent theory of consciousness^14,38,39^.

### Loss of consciousness prevents feature binding but not feature encoding

We further evaluated the effects of conscious states on the responses to deviant stimuli. In the awake state, participants exhibited robust responses to all types of deviant (Fig. 4A, top). Responses to deviants were stronger than those to the standards and yielded prominent mismatch responses. Temporal cluster-based permutation tests revealed that loudness deviant (LD) elicited earliest divergence (peak latency of 169 ms in a significant cluster ranging from 7 to 324 ms, *p* = 0.016), whereas tone deviant (TD) induced a later deviation from the responses to standard (peak latency of 195 ms in a significant cluster ranging from 116 to 400 ms, *p* = 0.020). Crucially, the combined deviant (CD) that contained the differences in both features evoked the strongest MMN response in a comparable latency as that in LD but earlier than that in TD (peak latency of 150 ms in a significant cluster ranging from 59 to 358 ms, *p* = 0.007). These results replicated the observations in scalp recordings^35^, demonstrating the feature integration in the auditory cortex with temporal-shift processing dynamics.

**Figure 4.**
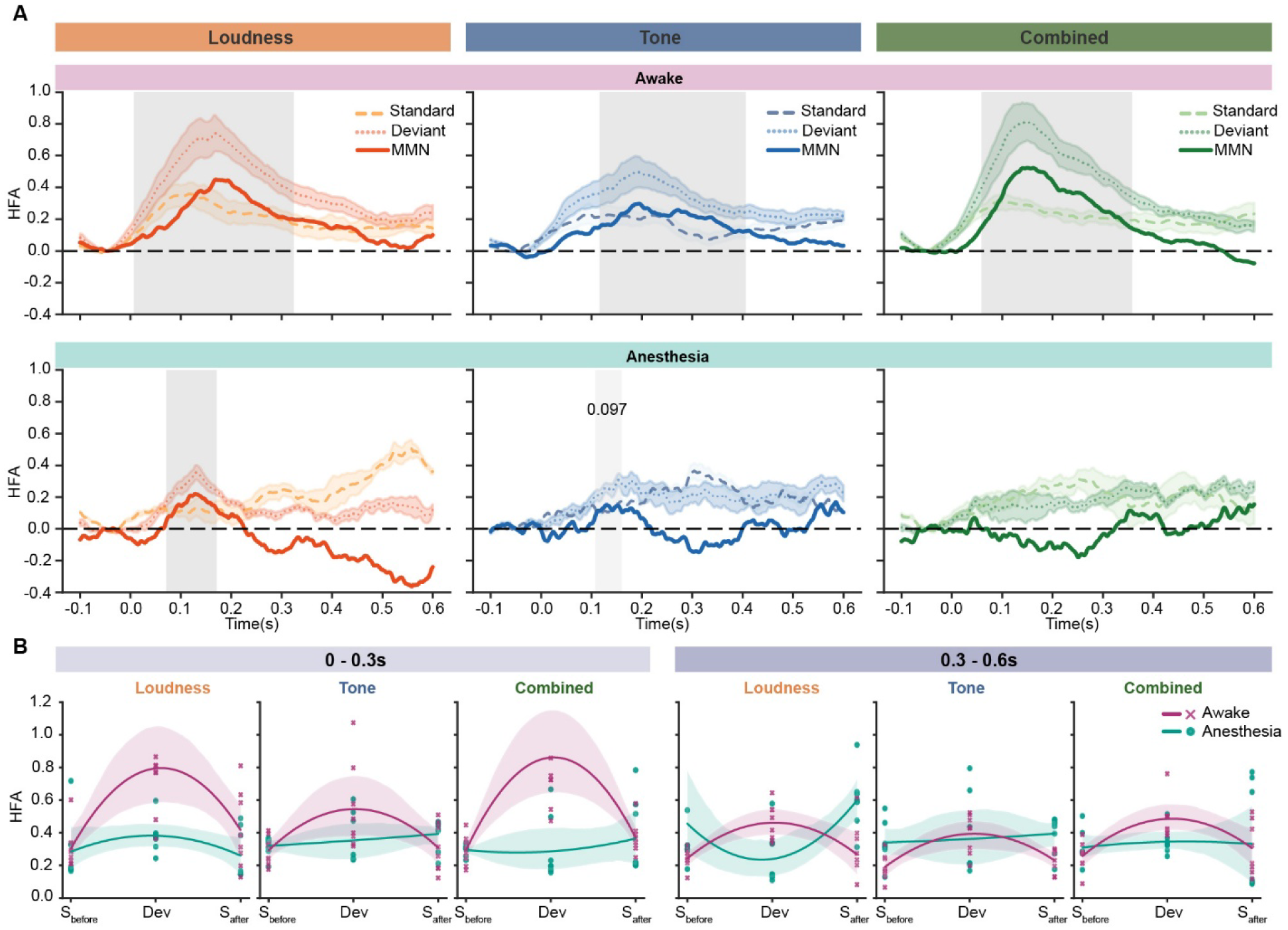
Loss of consciousness prevents feature binding but not encoding. A) Temporal profiles of mismatch responses in awake and anesthetized states. In the awake state (top), MMN responses were observed in each condition, as the significant temporal clusters were highlighted in shaded periods. In the anesthetized state (bottom), significant but shorter MMN responses were only in the single-feature deviant conditions, whereas no significant MMN was observed in the combined condition. B) Evolution of responses to the changes in features. In the early window (left, 0 to 0.3s), deviants (Dev) in the conscious state elicited stronger responses compared to the standards that are adjacent to the deviant; responses to the standards that are immediately before and after the deviant (S-before and S-after) evoked a similar level of responses. Whereas, in the anesthetic state, the deviant in the combined condition did not evoke different responses compared with adjacent standards. In the late window (right, 0.3 to 0.6s), the differences between the deviant and the adjacent standards were preserved in the awake state. In the anesthetized state, the response magnitude was comparable to that in the awake state, but the deviants did not show greater responses compared to adjacent standards. These results suggest that feature binding and local recurrent processes depend on consciousness.

In general anesthesia, a dissociation emerged between processes of single and combined attributes (Fig. 4A, bottom). Mismatch responses to the loudness deviant were observed (peak latency of 128 ms in a significant cluster ranging from 71 to 171 ms, *p* = 0.035); the mismatch responses to the tone deviant emerged with a weaker effect (peak latency of 130 ms in a cluster ranging from 108 to 160ms, *p* = 0.098). The duration of MMN in the anesthesized state was shorter compared to the sustained effects in the awake state, consistent with the local recurrent theory of consciousness^14,38,39^ and the loss of repetition effects to standard stimuli in a later process stage (Fig. 3). More importantly, the mismatch responses in the combined condition were absent. The deviant that included simultaneous changes of loudness and tone did not evoke any distinguishable responses compared to those of the standard stimuli. The absence of MMN to the combined deviant, contrasting with the observed MMN responses to the single feature deviants, suggests that anesthesia abolishes the automatic feature integration.

To further quantify the evolution of neural dynamics to feature change, we compared HFA elicited by the deviant stimuli against the responses to the standard stimuli that immediately preceded and followed the deviant using the ordinary least-squares (OLS) model. In the awake state, deviant stimuli evoked a clear enhancement in high-gamma activity relative to adjacent standards (Fig. 4B, left). The OLS model explained substantial variance in the early window (0–300 ms; R^2^ = 0.475, F(17,126) = 6.714, p < 0.001), revealing a pronounced quadratic, inverted U-shaped profile across trial positions (S_before_, Dev, S_after_), with the largest response at the deviant (β_quad = 0.177, *t*(126) = 6.544, p < 0.001) and no linear asymmetry was detected between pre- and post-deviant standards (β_linear = 0.039, p = 0.404). Feature comparisons revealed that the early deviance effect was strongest for combined deviants, slightly weaker for loudness, and weakest for lexical tone (difference between tone and combined: β = −0.0947, *t*(126) = −2.482, p = 0.014; loudness vs. combined: β = −0.0315, *t*(126) = −0.825, p = 0.411). This graded pattern indicates that early auditory mismatch responses scale with the complexity of feature change, consistent with early feature integration during wakefulness. Whereas, under anesthesia, both the overall high-gamma response and the deviant-centered inverted U-shape were markedly reduced (main state effect: β = −0.192, *t*(126) = −3.563, p < 0.001; state × quadratic: β = −0.191, *t*(126) = −5.009, p < 0.001). The mismatch pattern was absent for combined (β = −0.015) and tone deviants (β = −0.001); only a residual effect persisted for loudness (β = 0.037). These results indicate that anesthesia abolishes the evolution of neural responses between standard and deviant when features are presented simultaneously, and only retains minimal sensitivity for deviants of basic auditory attributes.

In the later window (300–600 ms, Fig. 4B, right), overall model fit declined (R^2^ = 0.285, F(17,126) = 2.955, p < 0.001), reflecting weaker and more variable late-stage activity. During wakefulness, a modest but reliable quadratic pattern persisted (β_quad = 0.0680, *t*(126) = 2.700, p = 0.008) in all conditions (quadratic × condition: F(2,126) = 2.418, p = 0.093). Under anesthesia, the response pattern changed (state × quadratic: F(2,126) = 21.81, p < 0.001). The quadratic effect was absent in combined (β = 0.008) and tone conditions (β = −0.002). The pattern was even reversed in the loudness condition (β = −0.0961), where responses to deviants were smaller than those of adjacent standards (three-way state × quadratic × loudness: β = −0.105, *t*(126) = −2.09, p = 0.038). These findings demonstrate that feedforward sensory mismatch computations for single features are qualitatively preserved in anesthesia, whereas the local recurrent computations that are necessary to sustain and integrate features depend on the conscious state.

### Consciousness orchestrates feature binding in a spatiotemporal hierarchy

To characterize the spatiotemporal architecture of automatic feature binding and the influence of the conscious state, we analyzed mismatch responses across anatomically distinct subregions of auditory cortex, pre-defined with functional localization in fMRI (Fig. 2), in both awake and anesthetized states. Given the vast differences in sites of implementation across individuals and the precise spatial requirements for testing specific hypotheses, we focused on two representative participants whose electrode implantation offers relatively consistent coverage of the auditory processing hierarchy—from anterior Heschl’s gyrus (aHG) through posterior HG (pHG) to mid and posterior superior temporal gyrus (mid-STG, pSTG). The unique spatial configuration enabled within-subject comparisons of neural dynamics across cortical fields, and the couple of available datasets were used for cross-validation^57^.

Participant sz01 had a coverage from pHG to pSTG (Fig. 5A). In the awake state, the averaged MMN amplitude across contacts was larger in the combined condition, compared to the two single attribute deviant conditions (Fig. 5B, left). The peak latency was earliest for the combined deviant (peak 148 ms), followed by loudness (183 ms), and tone (191 ms). The individual results were consistent with the group results in Fig. 4.

**Figure 5.**
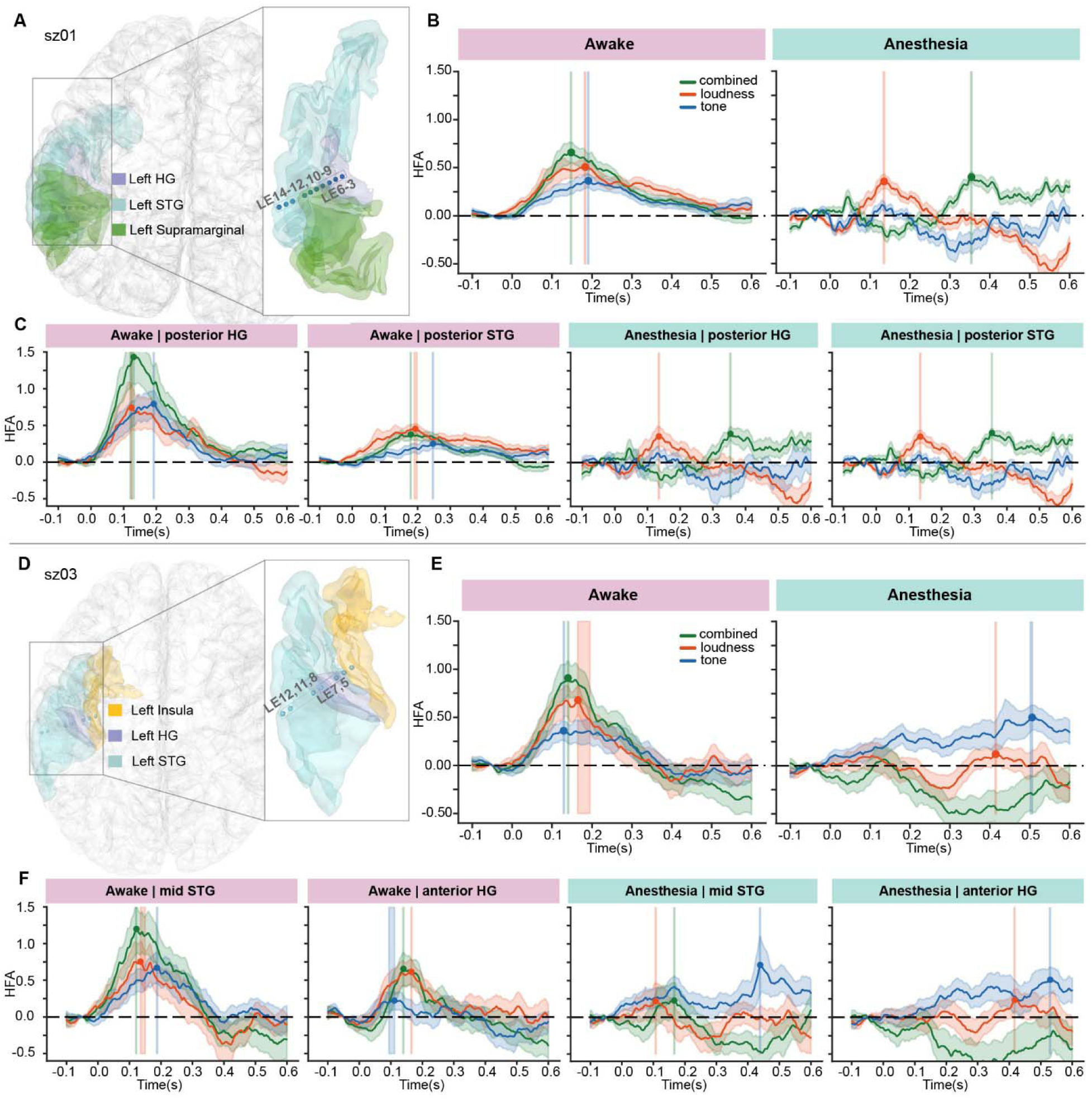
Consciousness orchestrates feature binding in a spatiotemporal hierarchy. A) Penetration of an electrode and anatomical location of contacts in a participant, sz01. The contacts covered the posterior Heschl’s gyrus (pHG) and posterior superior temporal gyrus (pSTG), which are regions for representing attributes of loudness and tone as identified in fMRI functional localization (Fig. 2). (B) MMN responses averaged across the contacts. In the awake state (left), combined deviants evoked stronger and faster responses compared with the deviants of single attributes of loudness and tone. In anesthetized state (right), only the loudness deviant elicited significant MMN in the early window; the responses in the combined condition were significantly delayed, with only comparable magnitude as that in the loudness condition. C) Distinct MMN responses in different auditory regions between conscious states. In the awake state (left, two plots), responses from contacts in pHG showed the characteristics of feature binding – stronger and faster MMN in combined conditions compared to those in single attribute deviant conditions Th location of pHG is consistent with the fMRI localization of overlapped representation of loudness and tone (Fig. 2). Whereas, responses in the combined condition from contacts in pSTG were comparable to those of single attribute deviant conditions. In the anesthetic state (right, two plots), the profiles of feature binding were absent in the same contacts from pHG and pSTG. D to F) Results for cross-validation in another participant, sz03. D) Penetration of an electrode and anatomical location of contacts in participant sz03. The contacts covered anterior Heschl’s gyrus (aHG) to middle superior temporal gyrus (mid-STG). E) Averaged MMN responses across the contacts. The results demonstrated similar characteristics of feature binding in the awake state and absence in the anesthetic state, as observed in B). F) Distinct MMN responses in different auditory regions between conscious states in sz03. The results qualitatively replicated the observations in C). Specifically, in the awake state, the profiles of feature binding were observed in mid-STG, which is another region of the co-representation of loudness and tone (Fig. 2); the profiles of binding were absent in aHG. Anesthesia eliminated the response profiles of feature binding in mid-STG and delayed the MMN responses into the late window. Peak MMN latencies were determined via non-parametric bootstrapping (1,000 resamples, 95% CI), indicated by bold points. Latency differences across conditions were considered significant when confidence intervals (vertical shaded areas around the latency points, ± SEM) did not overlap across conditions.

Whereas the response profile was different in the anesthesized state (Fig. 5B, right). In the early window, only the MMN responses for loudness were observed. The peak of MMN responses for the combined condition was postponed to the later window. The delayed responses were consistent with the observation that loss of consciousness influenced the later processing window (Figs 3&4). Moreover, the amplitude of the MMN responses in loudness and combined conditions was comparable, indicating no feature integration. These temporal and amplitude distinctions between awake and anesthesia suggest that loss of consciousness prevents automatic feature binding in the auditory cortex, consistent with the group results in Fig. 4.

Further dissecting according to the location of contacts, in the contacts of pHG (Fig. 5C, left) where the overlapped feature representation of loudness and tone existed (Fig. 2), the characteristics of feature binding were observed. Specifically, the MMN response amplitude was significantly larger than that of single attributes, as well as the peak latency of the combined condition was aligned with the earliest peak of loudness. Whereas in the contacts of pSTG where no overlapped representation of features was observed (Fig. 2), the profile of integration was absent – the MMN amplitude was comparable across three conditions. These results demonstrate that the automatic feature binding operates in a spatially organized auditory hierarchy.

However, in the anesthetized state, all the characteristics of feature binding were gone. Specifically, in the pHG and pSTG contacts, the peak of MMN responses in the combined condition was delayed into the late window after stimulus offset. Moreover, the amplitude difference across conditions observed in pHG in the awake state was eliminated, and showed comparable response magnitude between the combined and loudness conditions. These results suggest that the conscious state constrains the temporal dynamics at the particular level of auditory hierarchy for automatic feature binding.

Another independent sample of participant sz03 replicated the results obtained in participant sz01 about how conscious states modulate automatic feature integration in the auditory hierarchy. The participant sz03 had a coverage spanning aHG to mid-STG (Fig. 5D), capturing both core and belt auditory fields. In the awake state, deviant stimuli elicited the strongest responses for the combined condition, intermediate for loudness, and weakest for tone. These patterns mirrored the group-level results observed in participant sz01 (Fig. 5B), confirming that multiple-feature changes elicited the most robust prediction-error responses during consciousness. And anesthesia disrupts the feature integration process, postponing the responses to loudness and tone features.

Further dissecting by cortical site, contacts in the mid-STG exhibited the strongest mismatch responses to combined deviants, smaller but significant responses to loudness and to tone, demonstrating clear signatures of feature binding. The combined deviant also evoked the earliest peak (122 ms), followed by loudness (135 ms) and delayed tone responses (187 ms). The results replicated the response profile in pHG (sz01), supporting the temporal-shift processing dynamics during multi-feature integration^35^. In contrast, in aHG, where no overlapping representations of loudness and tone features were observed (Fig. 2), the integrative profile was absent. Combined deviants did not elicit an MMN response, and both single-feature responses were weak and delayed. This location of the observed effects, pHG in sz01 and mid-STG in sz03, aligns with the fMRI-decoded co-representation regions (Fig. 2), indicating that automatic feature binding is organized hierarchically in the auditory cortex.

In the anesthetized state, mismatch responses were substantially reduced and temporally varied (Fig. 5F, green traces). Specifically, in the mid-STG, the responses to combined deviants were reduced to the level of loudness responses, suggesting that feature-binding mechanisms were abolished. The consistent spatiotemporal patterns across both participants indicate that the conscious state constrains automatic feature binding across temporal and spatial hierarchies within local sensory networks.

## DISCUSSION

The present study probed the functional boundary of consciousness by investigating whether the conscious state is necessary for the integration of basic sensory features. By combining a novel multi-feature auditory oddball paradigm with intracranial sEEG recordings across awake and anesthetized states, we examined how the loss of consciousness alters feature encoding and binding within the auditory cortex. In the awake state, both feature encoding of single auditory attributes (tone and loudness) and cross-feature binding were automatically processed in localized sensory networks without requiring attention^27,58^. Under general anesthesia, the encoding of individual features remained intact, whereas feature binding in the integrative regions in the auditory cortices was selectively abolished^11,22^. These results reveal that the functional boundary of consciousness lies between the encoding and integration of basic sensory features, where the influences on the conscious-dependent computations^14,59^ are within both feedforward and recurrent processes in the local sensory neural network.

The observations of mismatch responses to single-feature deviants under anesthesia (Fig. 4A) are consistent with previous findings that basic sensory encoding can occur independently of awareness^27,60^. Automatic mismatch responses and early cortical activations are maintained in anesthetized animals and unconscious humans^33,37^, reflecting a feedforward, preconscious process of novelty detection. Our results support that the bottom-up encoding of sensory features and attributes does not rely on conscious access.

The absence of multi-feature mismatch responses in an anesthetized state (Fig. 4&5) functionally defines the lowest operational boundary of consciousness. The selective breakdown is at the level where independent auditory features are integrated into a unified perceptual representation^6,61,62^. In contrast to the existence of single feature encoding in anesthesia, feature binding collapses with the loss of consciousness, which marks the transition point between unconscious sensory registration and conscious perceptual construction. The dissociation provides direct neural evidence suggesting that binding, rather than basic encoding, is the first cortical computation that depends on consciousness^18,26^, and indicates the minimal neural mechanism that relates to conscious awareness.

Our results further reveal that the functional collapse occurs at an early feedforward stage of integration (Fig. 5). In the awake state, tone and loudness information converged rapidly in the posterior auditory regions – a candidate binding hub where co-representation for both attributes is identified by fMRI (Fig. 2). The short latency and localization of binding indicate that feature integration is initiated locally within the sensory hierarchy, but not at higher associative levels^1,63^. Under anesthesia, the feature integration vanished, demonstrating that feedforward binding is constrained by the conscious state. Previous findings have demonstrated that perceptual grouping and auditory object formation depend on conscious awareness ^18,59,64^; Our results push the temporal and functional limits forward by showing that even the initial feedforward convergence of features is disrupted when consciousness is lost. Thus, consciousness exerts its influence not only on higher-order cognitive synthesis but also on the earliest feedforward stage of feature integration.

In addition to the disruption of the feedforward process, our temporal analyses revealed that sustained responses were strongly influenced by anesthesia (Fig. 3D & 4B & 5). In the awake state, suppression in a later time window and prolonged mismatch responses were observed, presumably reflecting recurrent excitatory feedback for maintaining information, potentially predictive codes^65,66^. When consciousness was abolished, the recurrent loop collapsed, arguably unmasking inhibitory dynamics in local cortical circuits^22,67^. The differences between conscious states in the late activity window demonstrate that consciousness is also for sustaining recurrent computations, which possibly mediate the coherent binding of features into a unified percept.

By isolating a specific computation that vanishes with the loss of consciousness, our study provides a mechanistic test for competing theories of consciousness. By revealing a collapse of the integrative process in the sensory-level feedforward and recurrent loop, our results indicate that the minimal influence of consciousness resides at the local sensory circuit, where individual features integrate and form a coherent neural representation of a perceptual object. These findings align with the theories of local computations, such as Recurrent Processing Theory^14^, which propose that consciousness arises when sensory information is recurrently integrated across distributed cortical populations in local circuits. On the contrary, our results are inconsistent with the theories that emphasize global properties, such as Global Workspace frameworks^11^ and Integrated Information Theory (IIT), which predict that the functional boundary of consciousness is in higher-order associate areas.

The current findings significantly extend the depth of empirical studies on the function of consciousness. Previous study demonstrated semantic and syntactic integration as neural markers predicting recovery from vegetative states^68^. Whereas these studies probed semantic composition and sentence-level processing, our results reveal a more elementary boundary of consciousness that thresholds the binding of basic sensory features. The automatic feature integration may represent a foundational-level neural marker^25,26^, and the multi-feature MMN paradigm is a potential novel tool for assessing and predicting recovery in disorders of consciousness.

In conclusion, consciousness constrains the transition from feature encoding to feature integration, defining a functional boundary in local cortical computations. The conscious brain automatically binds sensory inputs into representations of unified perceptual objects, whereas loss of consciousness cuts off the integrative process. Our results identify the minimal functional boundary of consciousness and provide a computationally explicit neural marker for disorders of consciousness.

## Supporting information

for supplemental Figures and Tables

## RESOURCE AVAILABILITY

### Lead contact

Further information and requests for resources and reagents should be directed to and will be fulfilled by the lead contact, Xing Tian.

### Materials availability

The current study has not generated any new material.

### Data and code availability

- All original code has been deposited at OSF at osf.io/2t3hn and is publicly available as of the date of publication.
- Data from the representative participant (sz02) of all experimental conditions and all supplementary data tables are included in the OSF repository (https://osf.io/2t3hn) for verifying reproducibility and ensuring transparency. Because of ethical restrictions, the full dataset cannot be publicly archived. Readers interested in accessing the full dataset may contact the lead author. Data access will be granted to named individuals in accordance with ethical guidelines for the reuse of clinical data, subject to the completion of a formal data-sharing agreement and institutional approval.

## ACKNOWLEDGMENTS

We thank Zhiyuan Xu, Jiaqiu Sun and Xikang Luo for their constructive advice on data analysis. This study was supported by the National Natural Science Foundation of China (32271101), Program of Introducing Talents of Discipline to Universities Base B16018, and NYU Shanghai Boost Fund to X.Tian; and Shenzhen Science and Technology Innovation Committee (JCYJ20240813141105007, JCYJ20220818100213029) to C.Y.

## AUTHOR CONTRIBUTIONS

Conceptualization, Z.H and X.Tian; methodology, Z.H., H.Z., Q.C. and X.Tian; Investigation, Z.H., H.Z., X.C., Y.W., S.L., and C.Y; writing—original draft, Z.H.; writing—review & editing, All authors; funding acquisition, X.Teng, P.C.M.W., C.Y. and X.Tian; supervision, Z.H., C.Y. and X.Tian.

## DECLARATION OF GENERATIVE AI AND AI-ASSISTED TECHNOLOGIES

During the preparation of this work, the authors used ChatGPT to improve the readability and language of the manuscript. After using this tool or service, the authors reviewed and edited the content as needed and take full responsibility for the content of the publication.

## STAR⍰METHODS

### KEY RESOURCES TABLE

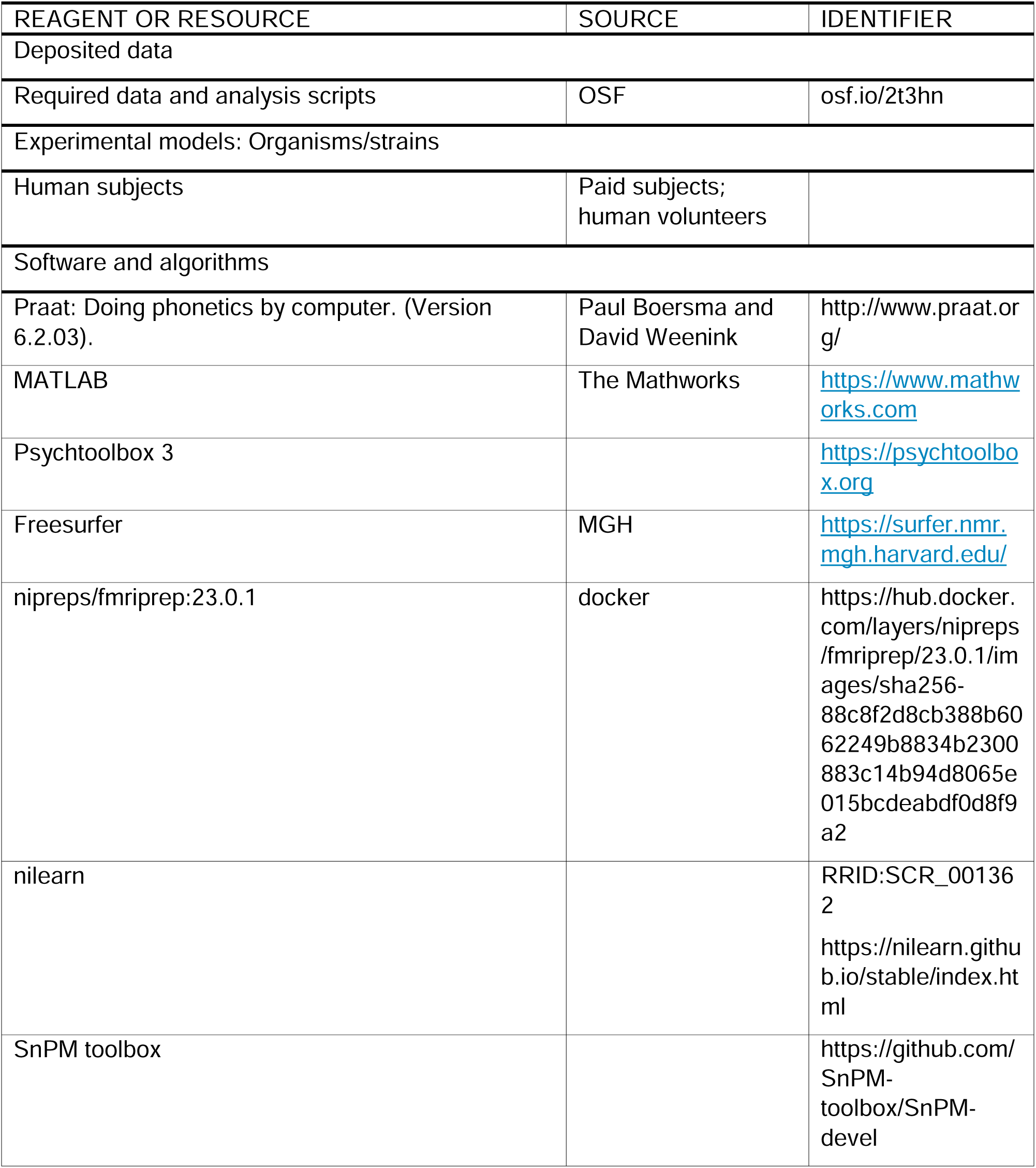
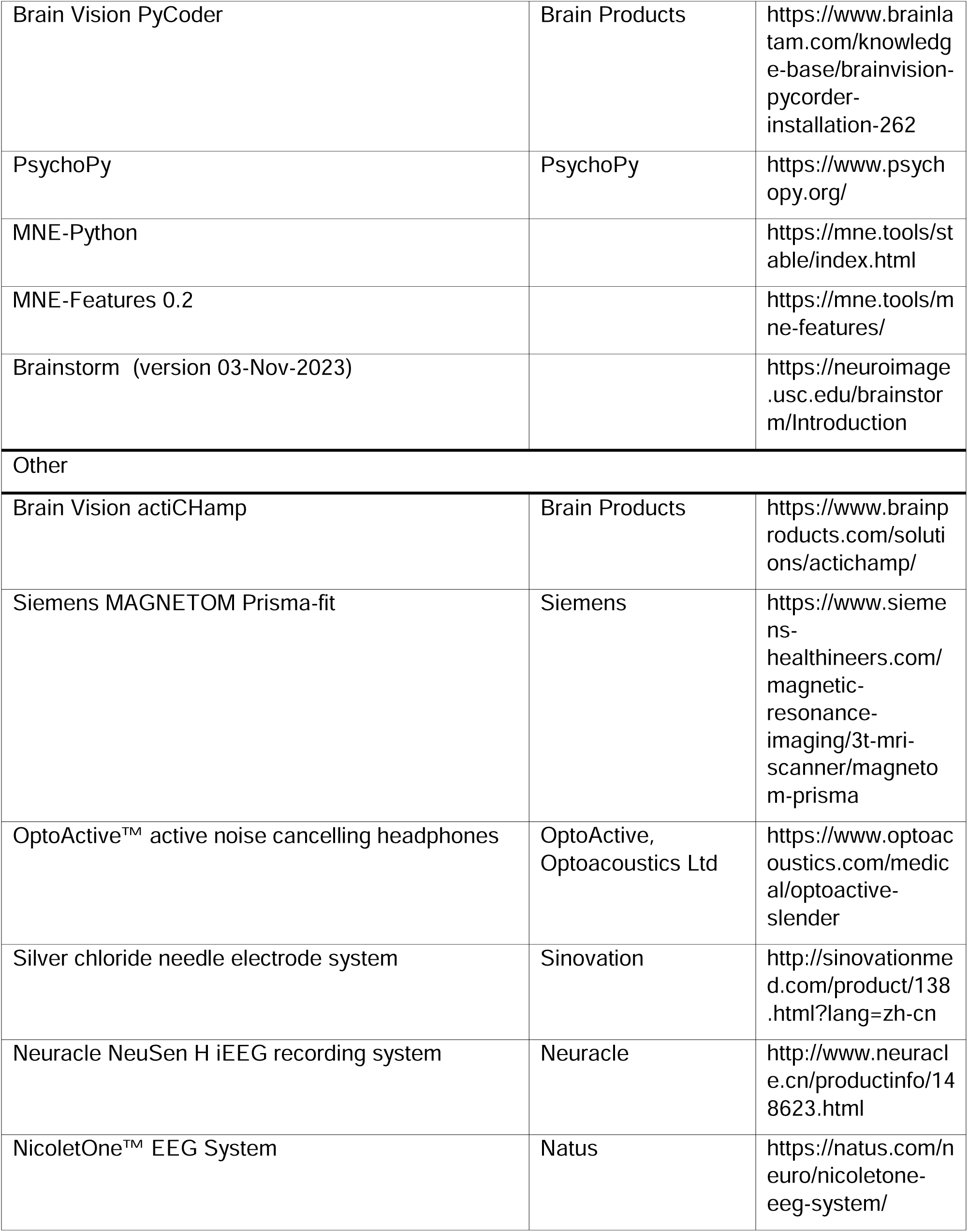
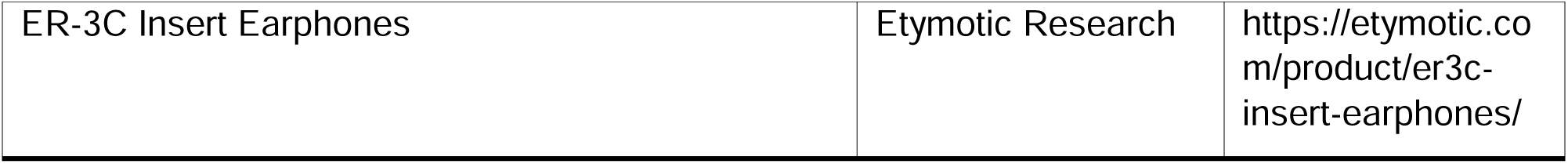

### EXPERIMENTAL MODEL AND STUDY PARTICIPANT DETAILS

#### fMRI experiment

fMRI data were acquired from 20 Mandarin-speaking volunteers (F = 16, M = 4), age between 19 to 26 years (Mean: 21.85). All participants are right-handed, with normal hearing (self-report) and no history of neurological or learning disorders. They received monetary incentives for their participation. The study was approved by the institutional review board at New York University Shanghai and conducted in accordance with the Declaration of Helsinki.

#### sEEG experiment

sEEG data were acquired from 13 Mandarin-speaking patients (F = 8, M = 5) who were undergoing clinical evaluation using sEEG for intractable epilepsy. Participants’ age ranges from 18 to 48 years old (Mean: 29) and they are all right-handed. All participants reported normal hearing. Clinically implanted depth electrodes (3.5 mm center-to-center spacing) were used to acquire sEEG data. A total of 2241 contacts (see Figure S1) were recorded from the 13 patients during the anesthesia and awake periods. During the anesthesia period, patients received a standardized intravenous regimen to ensure stable deep anesthesia. Anesthesia was induced with etomidate (20□mg, i.v.), sufentanil (20□μg, i.v.), and rocuronium bromide (50□mg, i.v.) to achieve rapid sedation, analgesia, and muscle relaxation. In addition, haloperidol (0.5□mg, i.v.) and dexamethasone (5□mg, i.v.) were administered to prevent agitation, nausea, and inflammation. Anesthesia was maintained using target-controlled infusion (TCI) of propofol and remifentanil via channels 2 and 3, respectively, with dosing parameters adjusted to maintain deep sedation without spontaneous movement or respiration. Dexmedetomidine was continuously infused at 0.2□μg/kg/h to provide additional sedation and hemodynamic stability. Supplemental oxygen mixed with medical air was delivered via mask at 2□L/min to ensure adequate oxygenation throughout the procedure. sEEG data were collected by performing an auditory oddball task during the deeply anesthetized state to ensure minimal behavioral or cognitive confounds. During the awake period, patients had a clear conversation with experimenters and followed instructions in an easy manner. The study was approved by the institutional review board at New York University Shanghai and Chen Zhen 2^nd^ People’s Hospital, and conducted in accordance with the Declaration of Helsinki.

### METHOD DETAILS

#### Stimuli and procedure

Mandarin tones were used as auditory stimuli^35^ in the perception task during fMRI experiment (Figure 1A top) and in the multi-feature oddball paradigm of the sEEG experiment across awake and anesthesia states (Figure 1A bottom). Specifically, Tone 1 and Tone 2 in Mandarin Chinese were recorded from a female native speaker in the way of humming. The stimuli were sampled at 44.1 kHz. The duration of the two stimuli was equalized to 300ms (see spectrograms in Figure S2).

##### fMRI experiment

We used Praat ^69^ to adjust the levels of intensity and created levels of loudness: 70 dB SPL (low loudness) and 85 dB SPL (high loudness). This calibration ensured that participants could clearly distinguish between soft and loud stimuli, as well as between Tone 1 and Tone 2 in the noisy environment of the MRI scanner. Combining the two tonal categories with the two loudness levels yielded four types of stimulus conditions (Fig. 1B, left): soft Tone 1 (sT1), loud Tone 1 (lT1), soft Tone 2 (sT2), and loud Tone 2 (lT2). Auditory stimuli were presented binaurally through MRI-compatible active noise-canceling headphones (OptoActive, Optoacoustics Ltd). Before headphone placement, participants also wore foam earplugs to further reduce scanner noise interference.

The experiment was presented and controlled by using MATLAB R2018b and the Psychophysics Toolbox ^70,71^. Before the main experiment, participants were familiarized with the four types of auditory stimuli. To simulate the acoustic environment of the actual scanning session, each sound stimulus was presented together with recorded MRI scanner noise. Participants reported the auditory stimuli to confirm their ability to identify each type of stimulus. This procedure ensured that participants could reliably discriminate among the stimulus types under continuous noise.

Inside the MRI scanner, participants performed the tone-listening task (Figure 1B). While lying supine in the scanner, participants viewed the experimental cues through a mirror mounted on the head coil. Each trial began with a fixation cross, time-locked to the onset of the auditory stimulus. Each mini-block consisted of five consecutive repetitions of the same auditory stimulus. The inter-stimulus interval (ISI) was dynamically adjusted based on stimulus duration and a variable delay, ensuring that each stimulus onset occurred exactly one second after the previous one. The repetition was designed to evoke fMRI adaptation effects ^72^. Stimuli in each mini-block were randomly selected from the four stimulus types (sT1, lT1, sT2, lT2), with each mini-block lasting 5 seconds and followed by a 10-second inter-block interval. A total of 20 mini-blocks were presented, comprising 100 stimulus trials in a run. To ensure that participants remained alert and were actively attending to the auditory stimuli, a ‘catch trial’ of a 1000 Hz pure tone was randomly inserted once in the first half (within the first 10 mini-blocks) and once in the second half (within the last 10 mini-blocks) of a run. Participants were instructed to press a button immediately upon detecting the target tone. Behavioral performance of the catch trials served as an index of participants’ attentiveness. The mini-blocks that included the catch trials were excluded from data analysis. After each run, participants had a rest and resumed the task after confirming their readiness. A total of 5 runs were included in the experiment, yielding a total of 100 mini-blocks and 450 analyzable trials.

##### sEEG experiment

The intensity of the auditory stimuli was modulated using Praat and yielded a low-loudness level (62 dB SPL) and a high-loudness level (81 dB SPL). By fully crossing tone and loudness levels, four types of auditory stimuli were generated: soft Tone 1 (sT1), loud Tone 1 (lT1), soft Tone 2 (sT2), and loud Tone 2 (lT2). Stimuli were delivered binaurally via air-conducted insert earphones (ER-3C Insert Earphones; Etymotic Research). Foam eartips of appropriate size were used to secure the earphones in the participants’ ear canals and prevent slippage.

The sEEG experiment was conducted while patients were anesthetized during clinical operation and awake during clinical monitoring in the wards. During surgery, patients did not have conscious awareness. Neurosurgeons played the sound sequence and recorded the neural responses. After surgery, patients sat upright in bed and listened to the same auditory stimuli as those presented intraoperatively. Participants were instructed to ignore the auditory stimuli and were free to engage in other activities (e.g., using a mobile phone or watching television).

A multi-feature oddball paradigm^35^ was used in the experiment (Fig. 1C). Standard and deviant stimuli were presented in a 4:1 ratio. To control for stimulus-specific effects, the experiment was divided into four sessions, with each session using one of the four auditory stimuli as the standard and the remaining three as deviants. This design yielded three deviant conditions: (1) Loudness Deviant (LD) – level of loudness changed compared with standard, (2) Tone Deviant (TD) – category of tone changed compared with standard, and (3) Combined Deviant (CD) – both attributes of loudness and tone changed compared with standard. The trial structure for each condition is summarized in Table S1. For example, in Session 1, the soft Tone 1 (sT1) stimulus was used as the standard (180 trials), and the remaining three stimuli served as deviants (15 trials each, totaling 45 deviant trials). Each session consisted of three sub-sessions targeting specific contrasts: (1) sT1 (60 standard trials) vs. lT1 (15 LD trials), (2) sT1 (60 standard trials) vs. sT2 (15 TD trials), and (3) sT1 (60 standard trials) vs. lT2 (15 CD trials). The presentation order of sub-sessions was pre-determined (Table S1). Within each sub-session, deviant stimuli were inserted in a pseudorandom order, with no deviant stimulus presented in consecutive trials nor placed at the beginning or end of a sequence. The duration of each sound and the inter-stimulus interval (ISI) were both set to 300ms. The entire experiment included 900 trials and lasted approximately 9 minutes.

#### Data acquisition

##### fMRI experiment

All MRI data, including structural and functional images, were acquired in the Child Brain Imaging Center (CBIC) at East China Normal University using a Siemens 3T MAGNETOM Prisma-fit MRI scanner. Before entering the scanner, participants were instructed to remain as still as possible during scanning. To minimize head motion artifacts, foam paddings were used to stabilize the head and secure the headphones in a 64-channel head coil.

High-resolution T1-weighted structural images were acquired using a magnetization-prepared rapid gradient echo (MPRAGE) sequence with the following parameters: repetition time (TR) = 2200 ms, echo time (TE) = 3.49 ms, flip angle = 8°, 192 slices, slice thickness = 1 mm, voxel size = 1×1×1 mm³, and field of view (FOV) = 256×256 mm².

Functional images were acquired using a T2*-weighted echo planar imaging (EPI) sequence with the following parameters: TR = 1000 ms, TE = 32 ms, flip angle = 55°, acceleration factor = 6, 72 slices, slice thickness = 2 mm, voxel size = 2×2×2 mm³, FOV = 192×192 mm², and interleaved slice acquisition. A total of five functional runs were collected, with each run consisting of 265 whole-brain volumes.

To correct for geometric distortions caused by magnetic field inhomogeneities—particularly prevalent in multiband EPI sequences—a field map was acquired before the functional runs. The field map consisted of two images with different echo times: a long TE (7.83 ms) and a short TE (4.92 ms), both with a TR of 413 ms. These field maps were used to correct for susceptibility-induced distortions, which typically affect regions such as the prefrontal cortex and mid-temporal areas, thereby reducing spatial signal loss in those regions.

##### sEEG experiment

Electrophysiological recordings were collected in the Department of Neurosurgery, Shenzhen Second People’s Hospital (First Affiliated Hospital of Shenzhen University). The signals were collected using two systems while patients were awake: (1) the Neuracle NeuSen H iEEG recording system, and (2) the NicoletOne™ EEG System. The first set of recording systems was mobile and was used during the surgical operation period while patients were anesthetized. Data was sampled at 1000 Hz and filtered online between 0 and 500 Hz. A white matter contact served as the online reference in both systems.

Intracranial depth electrodes were manufactured by Beijing Sinovation Medical Technology Co., Ltd. Each electrode contact had a diameter of 0.8 mm, a length of 2 mm, and an inter-contact spacing of 3.5 mm. The number of contacts per electrode ranged from 8 to 20. Before surgery, all participants underwent T1-weighted structural MRI scans to assess individual brain anatomy. After surgery, computed tomography (CT) scans were conducted to verify the location of electrode implantation. Data collection during the awake phase took place as part of routine postoperative electrophysiological monitoring, and recordings were only initiated when participants were in good physical condition, fully conscious, and had experienced no epileptic seizures within the preceding four hours.

#### Data pre-processing and artifact-rejection

##### fMRI experiment

Before preprocessing, all DICOM-format MRI files were organized according to the standardized Brain Imaging Data Structure (BIDS) format ^73^ using the *heudiconv* toolkit ^74^. During this step, data files were converted to NIfTI format, and each dataset was accompanied by a corresponding JSON sidecar file containing metadata about scanning parameters and acquisition details. The resulting BIDS-compliant dataset served as the input for subsequent data processing.

Preprocessing was performed using *fMRIPrep* ^75^, a robust and standardized pipeline designed to ensure reproducibility and transparency in fMRI analyses. *fMRIPrep* integrates tools from multiple widely adopted neuroimaging software packages—including AFNI ^76^, FSL ^77^, ANTs ^78^, FreeSurfer ^79^, and Nipype ^80^—and automatically selects the most optimal method at each processing step. For example, ANTs was used for spatial normalization, and FreeSurfer was employed for cortical surface reconstruction. This standardized approach enhances the reproducibility and reliability of the processed data.

To ensure full transparency and reproducibility of the processing workflow, all analyses were conducted using the *fMRIPrep* Docker container (version 23.0.1) executed on a Windows-based computing platform.

##### sEEG experiment

Localization of the stereo-electroencephalography (sEEG) electrode was performed using the Brainstorm toolbox ^81^ in combination with FreeSurfer ^79^, operating on a Linux (Ubuntu 22.04.3 LTS) platform with MATLAB R2023b. Preoperative T1-weighted MRI and postoperative CT scans were obtained from patients undergoing depth electrode implantation for clinical monitoring.

First, DICOM files were converted to NIfTI format using dcm2niix (v1.0.20220505) and organized according to the Brain Imaging Data Structure format ^73^. Cortical surface reconstruction was performed using FreeSurfer’s recon-all pipeline, generating anatomical surfaces (pial, white matter, mid-cortex) and volumetric parcellations (e.g., aseg.mgz, Destrieux atlas). These anatomical files were imported into Brainstorm to serve as the anatomical basis for electrode localization. Postoperative CT scans were co-registered to preoperative MRIs using affine transformation with ANTs ^78^ and visually inspected for alignment accuracy. The T1-weighted MRI served as the reference modality for subsequent normalization and atlas-based labeling. MNI ICBM152 nonlinear normalization was computed within Brainstorm to enable group-level spatial inference.

After co-registration and normalization, electrode trajectories were identified manually on co-registered CT images using Brainstorm’s sEEG/ECoG implantation module. Each electrode’s tip and skull entry point were annotated based on high-resolution CT slices, aided by supplementary electrode trajectory documentation provided by the clinical team. Naming conventions were standardized to ensure compatibility with Brainstorm (e.g., apostrophes and numerals were reformatted). Electrode models (Huake-Hengsheng SDE-08-S16) and contact counts were specified according to manufacturer specifications and implantation records. Electrode contact coordinates were projected onto two anatomical reference spaces: (1) Native space labeling, which is based on individual anatomy using the Destrieux cortical atlas; (2) MNI space labeling, in which contact locations were transformed to MNI space and labeled using the AAL3 atlas for cross-subject comparison. Both sets of anatomical labels were cross-referenced with electrophysiological recordings to validate anatomical targeting.

Electrode localization results were visualized within Brainstorm’s 3D cortical viewer to check whether the contacts were correctly located within the brain area. Cortex transparency, anatomical contours, and electrode positions were adjusted for visual clarity. For group-level and individual-level display, contact coordinates in MNI space were exported and rendered using MNE-Python^82^, enabling high-resolution interactive 3D visualization. FreeSurfer’s fsaverage brain was used as the template for surface projection.

sEEG data were preprocessed using a custom pipeline implemented in Python 3.12.4 with the MNE-Python library (v1.8.0), NumPy (v1.26.4), SciPy (v1.13.1), and other scientific computing libraries. Channels/contacts identified as epileptic or non-neural (including the trigger channel) were excluded. Additional channels dominated by 50 Hz line noise were detected using a custom notch-filtering and power thresholding approach and excluded if their post-filter energy exceeded the median plus ten times the mean absolute deviation. To minimize reference-dependent noise, data were re-referenced using the common average method.

A filtering process was applied to extract high gamma power, with a zero-phase bandpass filter (70–150 Hz) and notch filters (50, 100, 150, 200 Hz). Long epochs were extracted from −2000 ms to +2000 ms relative to stimulus onset to avoid edge effect, with baseline correction and detrending. Artifacts were identified via both peak-threshold and standard deviation criteria, flagging epochs with widespread high-amplitude fluctuations. Epochs were also reviewed manually using MNE’s interactive viewer.

### QUANTIFICATION AND STATISTICAL ANALYSIS

#### fMRI experiment

We first conducted univariate analysis to examine the spatial distribution of brain regions that represent acoustic attributes of loudness and tone. This method evaluates changes in BOLD signal at the voxel level to determine whether individual voxels show activity correlated with specific cognitive processes. Following a mini-block design (as described in the experimental procedures), repeated presentations of the same auditory stimulus within a mini-block were used to induce fMRI adaptation. Repetition-induced attenuation in BOLD responses serves as an index of feature invariance, i.e., selectivity to either tone or loudness^83,84^. Each mini-block was followed by a 10-second silent period to allow the BOLD responses to return to baseline, optimizing signal contrast. This design yields a predicted time series based on stimulus onset, duration, and event type, convolved with a canonical hemodynamic response function (HRF). The resulting model time series is compared to the measured BOLD signal at each voxel. A voxel is considered responsive to a condition if the model explains more variance than would be expected by chance. Correlation values from each voxel were aggregated into a whole-brain statistical map, with response magnitudes expressed in terms of effect size or significance levels. All analyses were conducted using the *nilearn* Python package ^85^.

For each participant, preprocessed structural and functional images were used in further analyses. At each voxel, the BOLD time series was modeled using a design matrix containing event regressors convolved with the double-gamma HRF from the SPM toolbox ^86^, including temporal derivatives to account for response variability. High-pass filtering (cutoff = 128 s) was applied, along with a cosine drift model to address low-frequency signal fluctuations and a first-order autoregressive model ^87^ to model serial autocorrelations and reduce noise.

Nuisance regressors from the preprocessing stage were included following denoising strategies common in resting-state fMRI ^88^ to enhance model fit without sacrificing temporal degrees of freedom. Each regressor and event contributed to a general linear model (GLM) fit to BOLD signals across five runs. The four auditory stimulus types (sT1, lT1, sT2, lT2) and their respective temporal derivatives were modeled as regressors of interest, with all other variables treated as nuisance regressors. Each mini-block (event) was defined by the onset of the first stimulus within the 5-second mini-block of five identical stimuli.

To identify regions selectively responding to tone and loudness, four contrasts were computed at the fixed-effects level: (1) general auditory activation (sT1:1, lT1:1, sT2:1, lT2:1), (2) loudness contrast (sT1:-1, lT1:1, sT2:-1, lT2:1), and (3) tone contrast (sT1:1, lT1:1, sT2:-1, lT2:-1). Each subject’s contrast maps were used in subsequent group-level analyses. To generalize across subjects, group-level analysis was performed using a random-effects model. One-sample t-tests were applied to each contrast, leveraging the fact that all functional data were normalized to MNI152NLin2009cAsym space during preprocessing, allowing voxel-wise comparison across participants. Each subject’s contrast map was spatially smoothed with an 8-mm FWHM Gaussian kernel, and non-grey matter voxels (e.g., white matter and CSF) were masked out. Statistical significance of the auditory contrast was determined using permutation-based nonparametric testing with 10,000 iterations and a corrected threshold of *p* < 0.05. Tone and loudness contrasts were thresholded at *p* < 0.01.

##### Multivariate analyses

Univariate analysis captures voxel-wise activation, but has limited power to distinguish differences spread across multiple voxels. Therefore, multivariate pattern analysis (MVPA) was employed, combining information from multiple voxels using machine learning classifiers to decode distributed representations ^89^. Integrating MVPA with repetition suppression provides a more sensitive approach to detecting distinct or overlapping neural representations^90^.

To implement MVPA, we used a searchlight approach ^91^, where a spherical region (radius = 10mm) centered on each voxel was used to extract multivoxel patterns for classification. Given the high dimensionality of fMRI data, this method avoids overfitting and memory issues while preserving local spatial information.

Classification was performed using Fisher’s Linear Discriminant Analysis (LDA), with shrinkage covariance estimation. T-values from the univariate GLM served as the response features. Classification accuracy for each sphere was computed, and chance-level accuracy was estimated using *scikit-learn*’s DummyClassifier ^92^. True accuracy was obtained by subtracting chance performance, and inter-subject pattern analysis (ISPA) was used to generalize decoding results across participants^93^.

##### ROI-based decoding analyses

Based on prior studies on pitch processing ^94,95^, we selected auditory-related ROIs from the DiFuMo 256-parcel atlas ^96^, including primary auditory cortex, temporoparietal junction, prefrontal cortex, and motor areas involved in speech production. ROI masks were created using nilearn and cross-referenced with Juelich, Destrieux, and Yeo atlases (see Table S2).

Labels were redefined for decoding loudness and tone: (1) for loudness decoding, sT1 and sT2 were labeled as soft; lT1 and lT2 as loud; (2) for tone decoding, sT1 and lT1 were labeled as Tone 1; sT2 and lT2 as Tone 2. This labeling ensured that irrelevant dimensions were ignored during decoding.

Data within each ROI were standardized and classified using LDA, with decision boundaries computed using least-squares fitting and regularization via the Ledoit-Wolf lemma. Decoding was repeated across the ROI using a 10-mm-radius searchlight. Classification results from 20 ROIs were validated via ISPA and statistically tested using nonparametric permutation testing with 1,000 iterations (SnPM toolbox), applying cluster-level correction (p < 0.05).

##### Preference Index Analysis

To assess feature selectivity, decoding results (after multiple comparison correction) were binarized (1 = significant, 0 = non-significant) to generate binary masks for subsequent analysis. Results were min-max normalized with the following equation, *x*′ = (*x* - *min*) / (*max* - *min*).

The *preference index*, *v*, was calculated with the following equation: *v* = *x*′ / (*x*′ - *x*′), where *x*′⍰ is the normalized loudness decoding score and x′⍰ is the normalized tone decoding score. The resulting *preference index, v,* ranges from 0 to 1, where 1 indicates a region prefers loudness and 0 indicates a preference for tone. This metric is independent of baseline signal intensity and is based on differences in representational strength across feature dimensions, providing a stable and interpretable index of feature selectivity.

#### sEEG experiment

##### Analysis of arousal level across conscious states

To first demonstrate the overall differences between anesthesia and awake states in intracranial recordings, we quantified the spectral slope of the power spectral density (PSD) of neural responses for each subject and state. Prior studies have demonstrated that the slope of the aperiodic 1/f component of cortical activity systematically steepens with decreasing levels of consciousness, such as during non-REM sleep and general anesthesia ^56^. This shift reflects a broadband reduction in high-frequency activity and has been proposed as a robust electrophysiological marker of arousal state.

For each subject and state (awake and anesthesia), preprocessed iEEG epochs were averaged per condition and subjected to power spectral analysis using Welch’s method. Power spectra were computed in the range of 1–150□Hz and log–log transformed to extract the aperiodic component. Spectral slope estimation was performed using the function from MNE-Features ^97^, which fits a linear model to the PSD in the log–log domain. The resulting slope and intercept parameters were averaged across all valid contacts or each patient.

This analysis served as a physiological control to verify that neural responses acquired during the anesthesia state were recorded in a state of diminished consciousness, thus providing an objective interpretive basis for subsequent comparisons of auditory processing across states.

##### Analysis of high-gamma response across conscious states

To investigate how cortical encoding of auditory stimuli differs between conscious and anesthetized states, we analyzed high-frequency activity (HFA, 70–150 Hz) extracted from sEEG recordings in auditory cortical electrodes. High-gamma power is a robust proxy for local neuronal population activity and has been widely used as a marker of feature-selective and perceptually relevant neural responses ^98,99^.

First, HFA amplitude was extracted using the Hilbert transform applied to band-pass filtered data (70–150Hz). The analytic amplitude was squared to yield power, log-transformed (in decibel scale), baseline-normalized using the pre-stimulus interval (−1000 to 0ms), and temporally smoothed with a 100ms moving average. Active auditory contacts were identified in the awake condition using paired-sample t-tests comparing mean HFA activity in the pre-stimulus (−100–0ms) and early post-stimulus (50–300ms) windows. Bonferroni correction was applied to control for multiple comparisons across contacts. Only contacts exhibiting significant HFA enhancement in the awake state were retained for further cross-state comparisons.

For each active contact, HFA responses were averaged across trials and extracted separately for the awake and anesthesia sessions. The resulting time series were compared using a non-parametric cluster-based permutation test ^100^ with 10,000 permutations, as implemented in MNE-Python. This test allowed us to identify time windows showing statistically significant differences in evoked HFA activity between states, while controlling for temporal dependencies.

All processing steps were implemented in a modular and reproducible Python pipeline using MNE-Python, and the full codebase has been shared on the Open Science Framework (OSF, osf.io/2t3hn) to support transparency and reuse.

##### Analysis of repetition-induced suppression for standard stimuli

To investigate whether suppression of repetitive auditory stimuli depends on conscious state, we analyzed HFA responses elicited by consecutively presented standard stimuli in the multi-feature oddball paradigm. This analysis compared responses across four sequential standard positions (S1 – S4) within each session (see Table 1 and Figure 1C).

We first restricted the analysis to contacts anatomically localized to the auditory cortex, which was defined by the significant auditory responses from the previous analysis using a combination of individual electrode labels and anatomical atlases (Desikan-Killiany and Destrieux). Contacts located in the Heschl’s gyrus (G_T_transv) and superior temporal gyrus (Plan_tempo, G_temp_sup-G_T_transv) were included. Anatomical localization was verified through MNI coordinates and visual inspection, and electrode names were standardized across patients. For the selected contacts, conditions (loudness, tone, combined), and states (awake, anesthesia), standard trials were subdivided according to their ordinal position (S1 through S4). Evoked responses for each standard position were averaged separately across trials, conditions and contacts within the auditory ROI. Outliers with extreme response amplitude (peak z-score > 3) were excluded to reduce the influence of transient artifacts. Further analysis was performed separately for two post-stimulus windows (0–0.3 s and 0.3–0.6 s) to investigate early and late processing components.

A two-way linear regression model was fit using ordinary least squares (OLS), with each standard’s ordinal position (S_position) treated as an ordinal predictor (1–4), and states were coded as a categorical factor for creating contrasts using the C function in statsmodels Python package, and Treatment for dummy coding:

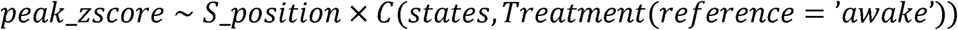

This analysis allowed us to test both main effects and the interaction between standard repetition position and conscious states. To visualize the response trajectories, the z-score trends were fitted using second-order polynomial fits across standard positions and states, which enables a direct comparison of repetition suppression dynamics across experimental conditions and states of consciousness.

##### Analysis of deviance-evoked mismatch responses across conscious states

Deviant-induced responses were extracted from contacts localized in the auditory cortex that showed reliable auditory-evoked activity, using the same preprocessing and selection criteria as described above. To assess the presence of deviance-related mismatch responses between three conditions across conscious states, we first performed a standard MMN analysis^27^, which is computed on high-gamma activity of deviant stimuli minus adjoining standard ones with a permutation test to specify the statistical significance.

Then we modeled the peak z-scored high-gamma amplitudes from the previous MMN responses using an OLS regression to further elucidate the response pattern underlying the MMN effect. That is, whether the enhanced deviant response arises from specific deviations relative to the adjacent standards. This analysis was performed separately for two post-stimulus windows (0–0.3 s and 0.3–0.6 s), corresponding to early and late processing components. For each contact, state (awake, anesthesia) and condition (LD, TD, CD) were defined as experimental factors. In each state and condition, three trial types were considered: the standard immediately preceding the deviant (*S_before_*), the deviant stimulus (*Dev*), and the standard immediately following the deviant (*S_after_*). Trials with extreme values (z > 3) were excluded.

To characterize the shape of deviance-related modulation, two orthogonal polynomial contrasts were constructed across trial type: (1) contrast_linear = (−1, 0, +1), capturing any asymmetry between pre- and post-deviant standards; (2) contrast_quad = (−1, +2, −1), capturing the quadratic “inverted U-shaped” profile corresponding to enhanced deviant responses relative to both adjacent standards.

The model was specified as:

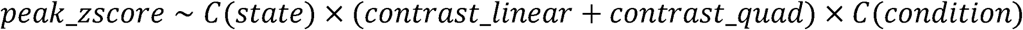

A significant positive quadratic contrast indicates that the deviant response is elevated relative to both flanking standards; the linear contrast reveals any directional asymmetry between pre- and post-deviant standards. Interactions with state and condition allowed us to assess how anesthesia modulates the strength and symmetry of deviance-related activity across different acoustic feature dimensions (tone, loudness, and combined). By fitting this model separately to the early (0–0.3 s) and late (0.3–0.6 s) windows, we further examined the temporal evolution of MMN-related activity and its sensitivity to consciousness level.

##### Analysis of Feature-Specific MMN Latency and Spatiotemporal Dynamics

To investigate the spatial and temporal architecture of auditory prediction errors, we quantified mismatch negativity (MMN) responses evoked by three types of deviants. The latency and magnitude of MMN responses were revealed in awake and anesthetized states across the auditory hierarchy.

Electrode contacts that were anatomically localized in ROIs of bilateral aHG, pHG, mid-STG, and pSTG were included. Moreover, only channels that showed significantly high-gamma activity to auditory stimuli in the awake state were included. Trial-wise MMN responses were computed as the difference between deviant and standard waveforms (–100 to 600 ms) for each condition. Peak latencies of MMN responses were estimated per region via bootstrapping. Specifically, within each combination of ROI × condition × state, difference waveforms were averaged across contacts in each ROI, and empirical peak latency (defined as time-to-max amplitude) was identified within the 0–600 ms post-stimulus window. To estimate the robustness of latency metrics, a non-parametric bootstrap procedure was implemented (1,000 resamples; confidence intervals, CIs at 95%). Thus, the 95% CI of latency estimates for each condition in each ROI was used to assess whether MMN peak latencies differed significantly across different conditions. Non-overlapping CIs between conditions were taken as evidence of significant temporal dissociation, indicating that different features (e.g., tone vs. loudness) elicited MMN responses at distinct latencies.

## REFERENCES

1. Bizley, J.K., and Cohen, Y.E. (2013). The what, where and how of auditory-object perception. Nat Rev Neurosci 14, 693–707. 10.1038/nrn3565.

2. Sussman, E.S. (2017). Auditory Scene Analysis: An Attention Perspective. J. Speech Lang. Hear. Res. 60, 2989–3000. 10.1044/2017_JSLHR-H-17-0041.

3. Jiang, Y., Zhou, K., and He, S. (2007). Human visual cortex responds to invisible chromatic flicker. Nat. Neurosci. 10, 657–662. 10.1038/nn1879.

4. Zou, J., He, S., and Zhang, P. (2016). Binocular rivalry from invisible patterns. Proc. Natl. Acad. Sci. 113, 8408–8413.

5. Treisman, A., and Gelade, G. (1980). A feature-integration theory of attention. Cognitive Psychology 12, 97–136. 10.1016/0010-0285(80)90005-5.

6. Treisman, A. (1999). Feature binding, attention and object perception. In Attention, space, and action: Studies in cognitive neuroscience (Oxford University Press), pp. 91–111.

7. Seth, A.K., and Bayne, T. (2022). Theories of consciousness. Nat Rev Neurosci 23, 439–452. 10.1038/s41583-022-00587-4.

8. Tononi, G., Boly, M., Massimini, M., and Koch, C. (2016). Integrated information theory: from consciousness to its physical substrate. Nat. Rev. Neurosci. 17, 450–461.

9. Albantakis, L., Barbosa, L., Findlay, G., Grasso, M., Haun, A.M., Marshall, W., Mayner, W.G.P., Zaeemzadeh, A., Boly, M., Juel, B.E., et al. (2023). Integrated information theory (IIT) 4.0: Formulating the properties of phenomenal existence in physical terms. PLOS Comput. Biol. 19, e1011465. 10.1371/journal.pcbi.1011465.

10. Baars, B.J. (2005). Global workspace theory of consciousness: toward a cognitive neuroscience of human experience. Prog. Brain Res. 150, 45–53.

11. Mashour, G.A., Roelfsema, P., Changeux, J.-P., and Dehaene, S. (2020). Conscious Processing and the Global Neuronal Workspace Hypothesis. Neuron 105, 776–798. 10.1016/j.neuron.2020.01.026.

12. Brown, R., Lau, H., and LeDoux, J.E. (2019). Understanding the higher-order approach to consciousness. Trends Cogn. Sci. 23, 754–768.

13. Lau, H., and Rosenthal, D. (2011). Empirical support for higher-order theories of conscious awareness. Trends Cogn. Sci. 15, 365–373.

14. Lamme, V.A.F. (2006). Towards a true neural stance on consciousness. Trends Cogn. Sci. 10, 494–501. 10.1016/j.tics.2006.09.001.

15. IIT-Concerned, Arnold, D.H., Baxter, M.G., Bekinschtein, T.A., Bengio, Y., Bisley, J.W., Browning, J., Buonomano, D., Carmel, D., Carrasco, M., et al. (2025). What makes a theory of consciousness unscientific? Nat. Neurosci. 28, 689–693. 10.1038/s41593-025-01881-x.

16. Frässle, S., Sommer, J., Jansen, A., Naber, M., and Einhäuser, W. (2014). Binocular rivalry: frontal activity relates to introspection and action but not to perception. J. Neurosci. 34, 1738–1747.

17. Cogitate Consortium, Ferrante, O., Gorska-Klimowska, U., Henin, S., Hirschhorn, R., Khalaf, A., Lepauvre, A., Liu, L., Richter, D., Vidal, Y., et al. (2025). Adversarial testing of global neuronal workspace and integrated information theories of consciousness. Nature 642, 133–142. 10.1038/s41586-025-08888-1.

18. Tsuchiya, N., Wilke, M., Frässle, S., and Lamme, V.A. (2015). No-report paradigms: extracting the true neural correlates of consciousness. Trends Cogn. Sci. 19, 757–770.

19. Owen, A.M., Coleman, M.R., Boly, M., Davis, M.H., Laureys, S., and Pickard, J.D. (2006). Detecting awareness in the vegetative state. Science 313, 1402–1402.

20. Dellert, T., Balster, H., Schlossmacher, I., Bruchmann, M., Moeck, R., and Straube, T. (2025). Neural correlates of consciousness in an auditory no-report fMRI study. Curr. Biol. 0, S096098222501334X. 10.1016/j.cub.2025.10.026.

21. Alkire, M.T., Hudetz, A.G., and Tononi, G. (2008). Consciousness and anesthesia. Science 322, 876–880.

22. Purdon, P.L., Pierce, E.T., Mukamel, E.A., Prerau, M.J., Walsh, J.L., Wong, K.F.K., Salazar-Gomez, A.F., Harrell, P.G., Sampson, A.L., Cimenser, A., et al. (2013). Electroencephalogram signatures of loss and recovery of consciousness from propofol. Proc. Natl. Acad. Sci. 110, E1142–E1151. 10.1073/pnas.1221180110.

23. Xiong, Y. (Sophy), Donoghue, J.A., Lundqvist, M., Mahnke, M., Major, A.J., Brown, E.N., Miller, E.K., and Bastos, A.M. (2024). Propofol-mediated loss of consciousness disrupts predictive routing and local field phase modulation of neural activity. Proc. Natl. Acad. Sci. 121, e2315160121. 10.1073/pnas.2315160121.

24. Schartner, M., Seth, A., Noirhomme, Q., Boly, M., Bruno, M.-A., Laureys, S., and Barrett, A. (2015). Complexity of Multi-Dimensional Spontaneous EEG Decreases during Propofol Induced General Anaesthesia. PLOS One 10, e0133532. 10.1371/journal.pone.0133532.

25. Casali, A.G., Gosseries, O., Rosanova, M., Boly, M., Sarasso, S., Casali, K.R., Casarotto, S., Bruno, M.-A., Laureys, S., Tononi, G., et al. (2013). A theoretically based index of consciousness independent of sensory processing and behavior. Sci. Transl. Med. 5, 198ra105. 10.1126/scitranslmed.3006294.

26. Mashour, G.A. (2024). Anesthesia and the neurobiology of consciousness. Neuron 112, 1553–1567. 10.1016/j.neuron.2024.03.002.

27. Näätänen, R., Paavilainen, P., Rinne, T., and Alho, K. (2007). The mismatch negativity (MMN) in basic research of central auditory processing: a review. Clinical neurophysiology 118, 2544–2590. https://doi.org/10/dd3b2j.

28. May, P.J.C., and Tiitinen, H. (2010). Mismatch negativity (MMN), the deviance-elicited auditory deflection, explained. Psychophysiology 47, 66–122. https://doi.org/10/bv4zsx.

29. Paavilainen, P. (2013). The mismatch-negativity (MMN) component of the auditory event-related potential to violations of abstract regularities: A review. International Journal of Psychophysiology 88, 109–123. https://doi.org/10/f429ng.

30. Paavilainen, P., Simola, J., Jaramillo, M., Näätänen, R., and Winkler, I. (2001). Preattentive extraction of abstract feature conjunctions from auditory stimulation as reflected by the mismatch negativity (MMN). Psychophysiology 38, 359–365. https://doi.org/10/dcqwxx.

31. Takegata, R., Brattico, E., Tervaniemi, M., Varyagina, O., Näätänen, R., and Winkler, I. (2005). Preattentive representation of feature conjunctions for concurrent spatially distributed auditory objects. Brain Res Cogn Brain Res 25, 169–179. https://doi.org/10/b2nfh8.

32. Sculthorpe, L.D., Ouellet, D.R., and Campbell, K.B. (2009). MMN elicitation during natural sleep to violations of an auditory pattern. Brain Res. 1290, 52–62.

33. Nourski, K.V., Steinschneider, M., Rhone, A.E., Kawasaki, H., Howard, M.A., and Banks, M.I. (2018). Auditory Predictive Coding across Awareness States under Anesthesia: An Intracranial Electrophysiology Study. J. Neurosci. 38, 8441–8452. 10.1523/JNEUROSCI.0967-18.2018.

34. Dykstra, A.R., and Gutschalk, A. (2015). Does the mismatch negativity operate on a consciously accessible memory trace? Sci Adv 1, e1500677. https://doi.org/10/gfw6vx.

35. Han, Z., Zhu, H., Shen, Y., and Tian, X. (2023). Segregation and integration of sensory features by flexible temporal characteristics of independent neural representations. Cerebral Cortex 33, 9542–9553.

36. Heinke, W., and Koelsch, S. (2005). The effects of anesthetics on brain activity and cognitive function. Curr. Opin. Anaesthesiol. 18, 625–631. 10.1097/01.aco.0000189879.67092.12.

37. Nourski, K.V., Steinschneider, M., Rhone, A.E., Kawasaki, H., Howard III, M.A., and Banks, M.I. (2018). Processing of auditory novelty across the cortical hierarchy: An intracranial electrophysiology study. Neuroimage 183, 412–424.

38. Lamme, V.A.F., and Roelfsema, P.R. (2000). The distinct modes of vision offered by feedforward and recurrent processing. Trends Neurosci. 23, 571–579. 10.1016/s0166-2236(00)01657-x.

39. Supèr, H., Spekreijse, H., and Lamme, V.A.F. (2001). Two distinct modes of sensory processing observed in monkey primary visual cortex (V1). Nat. Neurosci. 4, 304–310. 10.1038/85170.

40. Shamma, S.A., Elhilali, M., and Micheyl, C. (2011). Temporal coherence and attention in auditory scene analysis. Trends in Neurosciences 34, 114–123. 10.1016/j.tins.2010.11.002.

41. Zion Golumbic, E.M., Ding, N., Bickel, S., Lakatos, P., Schevon, C.A., McKhann, G.M., Goodman, R.R., Emerson, R., Mehta, A.D., Simon, J.Z., et al. (2013). Mechanisms Underlying Selective Neuronal Tracking of Attended Speech at a ‘Cocktail Party.’ Neuron 77, 980–991. 10.1016/j.neuron.2012.12.037.

42. Fries, P. (2015). Rhythms for Cognition: Communication through Coherence. Neuron 88, 220–235. 10.1016/j.neuron.2015.09.034.

43. Singer, W. (1999). Neuronal Synchrony: A Versatile Code for the Definition of Relations? Neuron 24, 49–65. 10.1016/S0896-6273(00)80821-1.

44. Engel, A.K., and Singer, W. (2001). Temporal binding and the neural correlates of sensory awareness. Trends in Cognitive Sciences 5, 16–25. 10.1016/S1364-6613(00)01568-0.

45. Uppenkamp, S., and Röhl, M. (2014). Human auditory neuroimaging of intensity and loudness. Hearing Research 307, 65–73. 10.1016/j.heares.2013.08.005.

46. Giraud, A.-L., Lorenzi, C., Ashburner, J., Wable, J., Johnsrude, I., Frackowiak, R., and Kleinschmidt, A. (2000). Representation of the temporal envelope of sounds in the human brain. J. Neurophysiol. 84, 1588–1598.

47. Hickok, G., and Poeppel, D. (2007). The cortical organization of speech processing. Nat Rev Neurosci 8, 393–402. https://doi.org/10/cpfwm5.

48. Si, X., Zhou, W., and Hong, B. (2017). Cooperative cortical network for categorical processing of Chinese lexical tone. Proc Natl Acad Sci USA 114, 12303–12308. https://doi.org/10/gcmwxg.

49. Liang, B., and Du, Y. (2018). The Functional Neuroanatomy of Lexical Tone Perception: An Activation Likelihood Estimation Meta-Analysis. Frontiers in Neuroscience 12, 495. https://doi.org/10/gd2gg6.

50. Rauschecker, J.P., and Scott, S.K. (2009). Maps and streams in the auditory cortex: nonhuman primates illuminate human speech processing. Nat Neurosci 12, 718–724. https://doi.org/10/bz55p6.

51. Liang, B., Li, Y., Zhao, W., and Du, Y. (2023). Bilateral human laryngeal motor cortex in perceptual decision of lexical tone and voicing of consonant. Nat. Commun. 14, 4710. 10.1038/s41467-023-40445-0.

52. Auksztulewicz, R., and Friston, K. (2016). Repetition suppression and its contextual determinants in predictive coding. Cortex 80, 125–140. 10.1016/j.cortex.2015.11.024.

53. Grill-Spector, K., Henson, R., and Martin, A. (2006). Repetition and the brain: neural models of stimulus-specific effects. Trends in Cognitive Sciences 10, 14–23. 10.1016/j.tics.2005.11.006.

54. Summerfield, C., Trittschuh, E.H., Monti, J.M., Mesulam, M.-M., and Egner, T. (2008). Neural repetition suppression reflects fulfilled perceptual expectations. Nat. Neurosci. 11, 1004–1006. 10.1038/nn.2163.

55. Vuilleumier, P., Henson, R.N., Driver, J., and Dolan, R.J. (2002). Multiple levels of visual object constancy revealed by event-related fMRI of repetition priming. Nat Neurosci 5, 491–499. 10.1038/nn839.

56. Lendner, J.D., Helfrich, R.F., Mander, B.A., Romundstad, L., Lin, J.J., Walker, M.P., Larsson, P.G., and Knight, R.T. (2020). An electrophysiological marker of arousal level in humans. eLife 9, e55092. 10.7554/eLife.55092.

57. Mercier, M.R., Dubarry, A.-S., Tadel, F., Avanzini, P., Axmacher, N., Cellier, D., Vecchio, M.D., Hamilton, L.S., Hermes, D., Kahana, M.J., et al. (2022). Advances in human intracranial electroencephalography research, guidelines and good practices. NeuroImage, 119438. 10.1016/j.neuroimage.2022.119438.

58. Garrido, M.I., Kilner, J.M., Stephan, K.E., and Friston, K.J. (2009). The mismatch negativity: A review of underlying mechanisms. Clinical Neurophysiology 120, 453–463. https://doi.org/10/bcv6fh.

59. Dehaene, S., and Changeux, J.-P. (2011). Experimental and Theoretical Approaches to Conscious Processing. Neuron 70, 200–227. 10.1016/j.neuron.2011.03.018.

60. Koch, C., Massimini, M., Boly, M., and Tononi, G. (2016). Neural correlates of consciousness: progress and problems. Nat. Rev. Neurosci. 17, 307–321.

61. Lamme, V.A.F. (2010). How neuroscience will change our view on consciousness. Cogn. Neurosci. 1, 204–220. 10.1080/17588921003731586.

62. Dehaene, S., Changeux, J.-P., Naccache, L., Sackur, J., and Sergent, C. (2006). Conscious, preconscious, and subliminal processing: a testable taxonomy. Trends Cogn. Sci. 10, 204–211. 10.1016/j.tics.2006.03.007.

63. Griffiths, T.D., and Warren, J.D. (2004). What is an auditory object? Nat. Rev. Neurosci. 5, 887–892. 10.1038/nrn1538.

64. Wacongne, C., Labyt, E., van Wassenhove, V., Bekinschtein, T., Naccache, L., and Dehaene, S. (2011). Evidence for a hierarchy of predictions and prediction errors in human cortex. Proceedings of the National Academy of Sciences 108, 20754–20759. https://doi.org/10/fqr69h.

65. Friston, K. (2005). A theory of cortical responses. Philos. Trans. R. Soc. B: Biol. Sci. 360, 815–836. 10.1098/rstb.2005.1622.

66. Bastos, A.M., Usrey, W.M., Adams, R.A., Mangun, G.R., Fries, P., and Friston, K.J. (2012). Canonical Microcircuits for Predictive Coding. Neuron 76, 695–711. 10.1016/j.neuron.2012.10.038.

67. Brown, E.N., Lydic, R., and Schiff, N.D. (2010). General Anesthesia, Sleep, and Coma. N. Engl. J. Med. 363, 2638–2650. 10.1056/NEJMra0808281.

68. Gui, P., Jiang, Y., Zang, D., Qi, Z., Tan, J., Tanigawa, H., Jiang, J., Wen, Y., Xu, L., Zhao, J., et al. (2020). Assessing the depth of language processing in patients with disorders of consciousness. Nat. Neurosci. 23, 761–770. 10.1038/s41593-020-0639-1.

69. Boersma, P., and Weenink, D. (2021). Praat: doing phonetics by computer. Version Version 6.2.03.

70. Brainard, D.H. (1997). The Psychophysics Toolbox. Spatial Vision 10, 433–436. 10.1163/156856897X00357.

71. Mario, K., Brainard, D., and Pelli, D. (2007). What’s new in Psychtoolbox-3?

72. Leaver, A.M., and Rauschecker, J.P. (2010). Cortical Representation of Natural Complex Sounds: Effects of Acoustic Features and Auditory Object Category. J. Neurosci. 30, 7604–7612. 10.1523/JNEUROSCI.0296-10.2010.

73. Gorgolewski, K.J., Auer, T., Calhoun, V.D., Craddock, R.C., Das, S., Duff, E.P., Flandin, G., Ghosh, S.S., Glatard, T., Halchenko, Y.O., et al. (2016). The brain imaging data structure, a format for organizing and describing outputs of neuroimaging experiments. Sci Data 3, 160044. 10.1038/sdata.2016.44.

74. Halchenko, Y., Goncalves, M., Velasco, P., Castello, M.V. di O., Ghosh, S., Salo, T., II, J.T.W., Hanke, M., Sadil, P., Christian, H., et al. (2023). nipy/heudiconv: v1.0.0. Version v1.0.0 (Zenodo).

75. Esteban, O., Markiewicz, C.J., Blair, R.W., Moodie, C.A., Isik, A.I., Erramuzpe, A., Kent, J.D., Goncalves, M., DuPre, E., Snyder, M., et al. (2019). fMRIPrep: a robust preprocessing pipeline for functional MRI. Nat Methods 16, 111–116. 10.1038/s41592-018-0235-4.

76. Cox, R.W. (1996). AFNI: Software for Analysis and Visualization of Functional Magnetic Resonance Neuroimages. Computers and Biomedical Research 29, 162–173. 10.1006/cbmr.1996.0014.

77. Jenkinson, M., Beckmann, C.F., Behrens, T.E.J., Woolrich, M.W., and Smith, S.M. (2012). FSL. NeuroImage 62, 782–790. 10.1016/j.neuroimage.2011.09.015.

78. Avants, B.B., Tustison, N., Song, G., and others (2009). Advanced normalization tools (ANTS). Insight j 2, 1–35.

79. Fischl, B. (2012). FreeSurfer. Neuroimage 62, 774–781.

80. Gorgolewski, K., Burns, C.D., Madison, C., Clark, D., Halchenko, Y.O., Waskom, M.L., and Ghosh, S.S. (2011). Nipype: a flexible, lightweight and extensible neuroimaging data processing framework in python. Frontiers in neuroinformatics 5, 13.

81. Tadel, F., Baillet, S., Mosher, J.C., Pantazis, D., and Leahy, R.M. (2011). Brainstorm: A User-Friendly Application for MEG/EEG Analysis. Comput. Intell. Neurosci. 2011, 1–13. 10.1155/2011/879716.

82. Gramfort, A. (2013). MEG and EEG data analysis with MNE-Python. Front. Neurosci. 7. https://doi.org/10/gdvhvz.

83. Grill-Spector, K., Kushnir, T., Edelman, S., Avidan, G., Itzchak, Y., and Malach, R. (1999). Differential Processing of Objects under Various Viewing Conditions in the Human Lateral Occipital Complex. Neuron 24, 187–203. 10.1016/S0896-6273(00)80832-6.

84. Kourtzi, Z., and Kanwisher, N. (2001). Representation of Perceived Object Shape by the Human Lateral Occipital Complex. Science 293, 1506–1509. 10.1126/science.1061133.

85. Nilearn contributors (2022). nilearn. 10.5281/zenodo.8397156 https://doi.org/10.5281/zenodo.8397156.

86. Penny, W.D., Friston, K.J., Ashburner, J.T., Kiebel, S.J., and Nichols, T.E. (2011). Statistical parametric mapping: the analysis of functional brain images (Elsevier).

87. Bullmore, E., Brammer, M., Williams, S.C., Rabe-Hesketh, S., Janot, N., David, A., Mellers, J., Howard, R., and Sham, P. (1996). Statistical methods of estimation and inference for functional MR image analysis. Magnetic Resonance in Medicine 35, 261–277.

88. Fox, M.D., Snyder, A.Z., Vincent, J.L., Corbetta, M., Van Essen, D.C., and Raichle, M.E. (2005). The human brain is intrinsically organized into dynamic, anticorrelated functional networks. Proceedings of the National Academy of Sciences 102, 9673–9678. 10.1073/pnas.0504136102.

89. Norman, K.A., Polyn, S.M., Detre, G.J., and Haxby, J.V. (2006). Beyond mind-reading: multi-voxel pattern analysis of fMRI data. Trends in Cognitive Sciences 10, 424–430. 10.1016/j.tics.2006.07.005.

90. Barron, H.C., Garvert, M.M., and Behrens, T.E.J. (2016). Repetition suppression: a means to index neural representations using BOLD? Philosophical Transactions of the Royal Society B: Biological Sciences 371, 20150355. 10.1098/rstb.2015.0355.

91. Kriegeskorte, N., Goebel, R., and Bandettini, P. (2006). Information-based functional brain mapping. Proceedings of the National Academy of Sciences 103, 3863–3868. 10.1073/pnas.0600244103.

92. Pedregosa, F., Varoquaux, G., Gramfort, A., Michel, V., Thirion, B., Grisel, O., Blondel, M., Prettenhofer, P., Weiss, R., Dubourg, V., et al. (2011). Scikit-learn: Machine learning in Python. the Journal of machine Learning research 12, 2825–2830.

93. Wang, Q., Cagna, B., Chaminade, T., and Takerkart, S. (2020). Inter-subject pattern analysis: A straightforward and powerful scheme for group-level MVPA. NeuroImage 204, 116205. 10.1016/j.neuroimage.2019.116205.

94. Czoschke, S., Fischer, C., Bahador, T., Bledowski, C., and Kaiser, J. (2021). Decoding Concurrent Representations of Pitch and Location in Auditory Working Memory. J. Neurosci. 41, 4658–4666. https://doi.org/10/gj7mv3.

95. May, L., Halpern, A.R., Paulsen, S.D., and Casey, M.A. (2022). Imagined Musical Scale Relationships Decoded from Auditory Cortex. Journal of Cognitive Neuroscience 34, 1326–1339. 10.1162/jocn_a_01858.

96. Dadi, K., Varoquaux, G., Machlouzarides-Shalit, A., Gorgolewski, K.J., Wassermann, D., Thirion, B., and Mensch, A. (2020). Fine-grain atlases of functional modes for fMRI analysis. NeuroImage 221, 117126. 10.1016/j.neuroimage.2020.117126.

97. Schiratti, J.-B., Le Douget, J.-E., Le Van Quyen, M., Essid, S., and Gramfort, A. (2018). An Ensemble Learning Approach to Detect Epileptic Seizures from Long Intracranial EEG Recordings. In 2018 IEEE International Conference on Acoustics, Speech and Signal Processing (ICASSP), pp. 856–860. 10.1109/ICASSP.2018.8461489.

98. Edwards, E., Soltani, M., Deouell, L.Y., Berger, M.S., and Knight, R.T. (2005). High Gamma Activity in Response to Deviant Auditory Stimuli Recorded Directly From Human Cortex. Journal of Neurophysiology 94, 4269–4280. 10.1152/jn.00324.2005.

99. Lachaux, J.-P., Axmacher, N., Mormann, F., Halgren, E., and Crone, N.E. (2012). High-frequency neural activity and human cognition: past, present and possible future of intracranial EEG research. Prog. Neurobiol. 98, 279–301. 10.1016/j.pneurobio.2012.06.008.

100. Maris, E., and Oostenveld, R. (2007). Nonparametric statistical testing of EEG-and MEG-data. Journal of Neuroscience Methods 164, 177–190. https://doi.org/10/dt923p.

