## Supplementary material for "Automatic binding of basic sensory features requires consciousness": for supplemental Figures and Tables

**SUPPLEMENTAL INFORMATION**

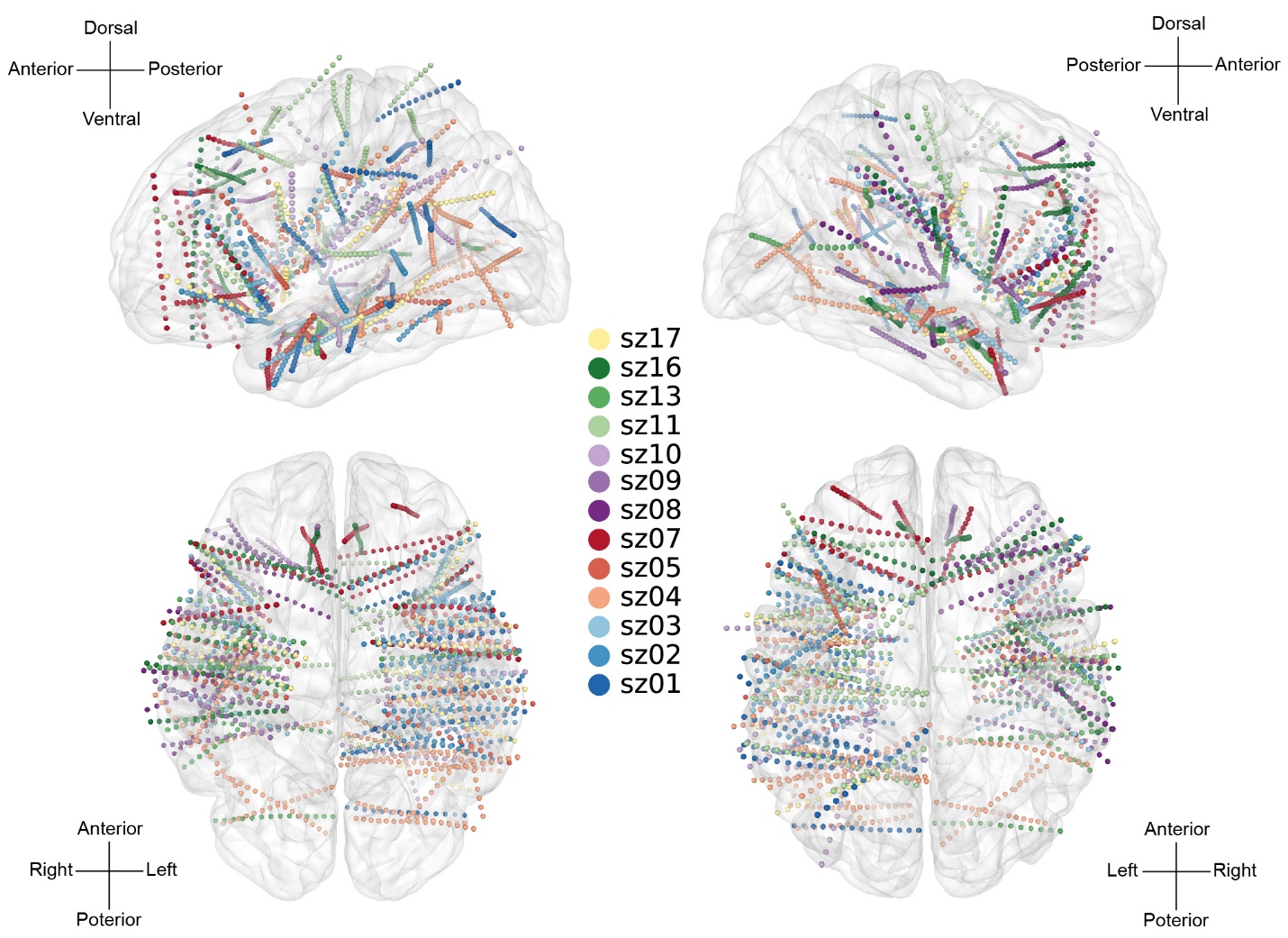

Figure S1. Contact localization from all patients.

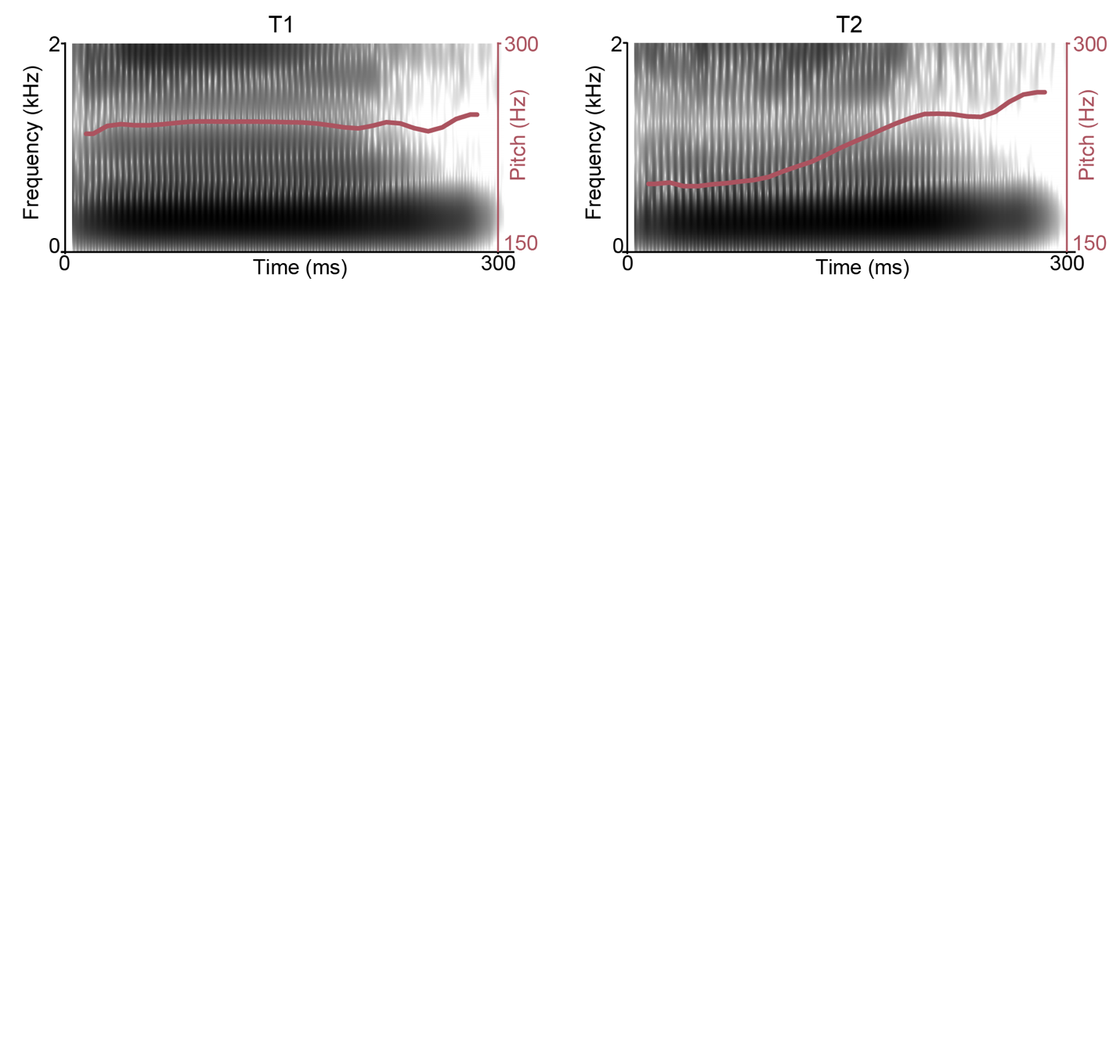

Figure S2 Spectrograms of the Mandarin tone stimuli used in the experiment. The left panel shows the spectrographic profile of a high-level tone (Tone 1), and the right panel shows a rising tone (Tone 2). Red lines superimposed on the spectrograms indicate the extracted fundamental frequency (F0) contours over time.

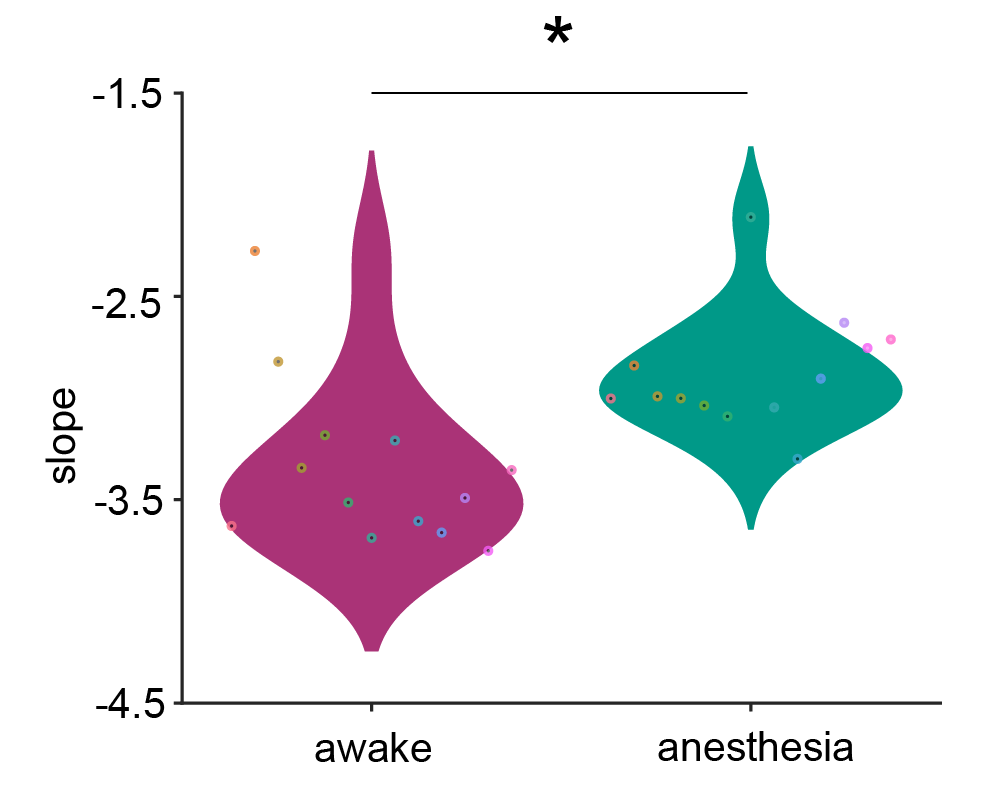

Figure S3 Quantification of arousal state using the spectral slope of the aperiodic 1/f power spectrum. Log–log power spectral density (PSD) was estimated from preprocessed sEEG recordings. Spectral slopes were significantly steeper during anesthesia than in the awake state (t = −3.13, p < 0.05, paired-sample t test), consistent with a suppression of high-frequency activity and diminished cortical excitability. This confirmed that intraoperative recordings were obtained during physiologically unconscious states. Spectral slope analysis was computed across all valid contacts per patient.

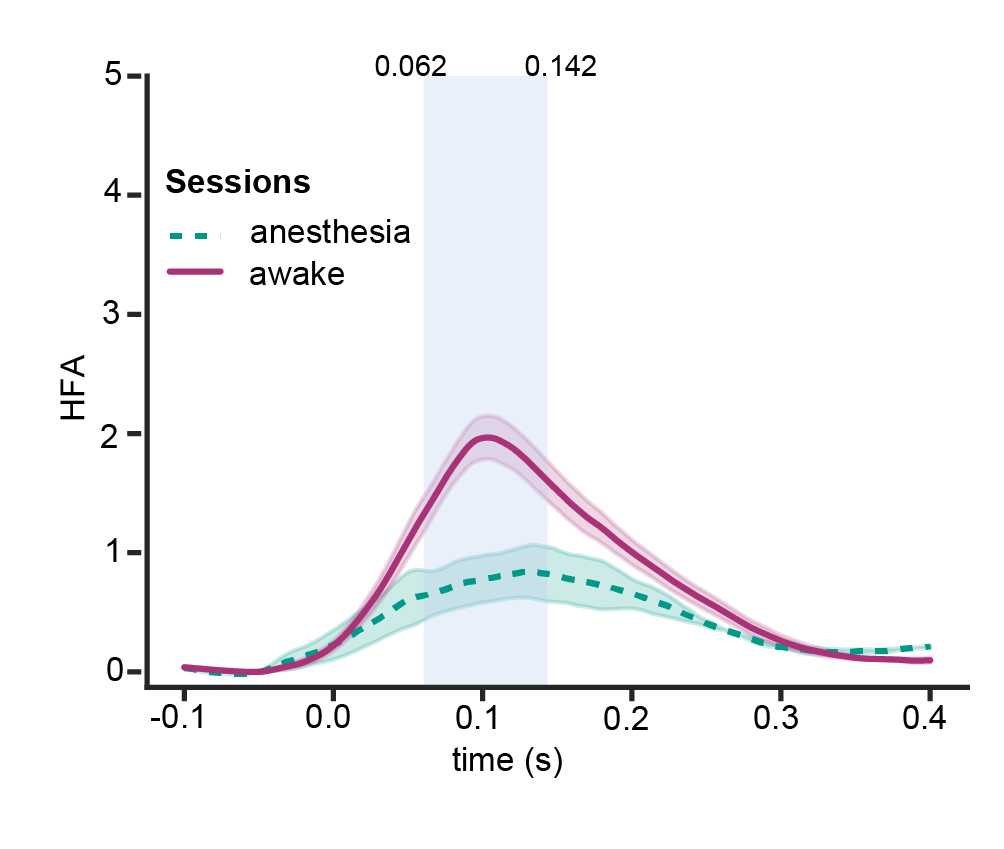

Figure S4 Group-level intracranial high-gamma responses (70–150 Hz). Significantly stronger responses were observed in the awake state relative to anesthesia, by a t-test on the selected auditory contacts (z > 1.65) in Fig. 3A. Significant temporal clusters were identified using non-parametric cluster-based permutation testing (10,000 permutations, p = 0.022; shaded region shows significant time window: 62–142 ms). Error bars represent ± SEM.

***Table S1. The conditions in the sEEG multi-feature deviant oddball paradigm. Numbers in parentheses indicate the trial number for the stimulus.***

| Stimulus types | Session 1 | Session 2 | Session 3 | Session 4 |
| --- | --- | --- | --- | --- |
| Standard | sT1 (180) | lT1 (180) | sT2 (180) | lT2(180) |
| Loudness deviant | lT1 (15) | sT1 (15) | lT2 (15) | sT2 (15) |
| Tone deviant | sT2 (15) | lT2 (15) | sT1 (15) | lT1 (15) |
| Combined deviant | lT2 (15) | sT2 (15) | lT1 (15) | sT1 (15) |

***Table S2. Cortical ROIs used for fMRI decoding in Juelich, Destrieux, and Yeo atlases.***

| **Destrieux** | **Juelich** | **Yeo et al. 20ll 17 network** |
| --- | --- | --- |
| L G_temp_sup-Plan_tempo | GM Inferior parietal lobule PFm L | SalVentAttnA |
| L S_temporal_sup | GM Inferior parietal lobule PFm L | TempPar |
| L S_interm_prim-Jensen | GM Inferior parietal lobule Pga L | DefaultB |
| R G_pariet_inf-Angular | GM Inferior parietal lobule PGp R | DefaultC |
| R G_pariet_inf-Angular | GM Inferior parietal lobule Pga R | DefaultA |
| R G_pariet_inf-Angular | GM Inferior parietal lobule Pga L | ContB |
| R G_pariet_inf-Angular | GM Inferior parietal lobule Pga R | ContB |
| L G_pariet_inf-Angular | GM Inferior parietal lobule PGp L | DefaultA |
| R S_interm_prim-Jensen | GM Inferior parietal lobule PFm R | TempPar |
| RS_central | GM Primary motor cortex BA4p L | SomIMotA |
| R G_postcentral | GM Primary somatosensory cortex BA1 R | SomIMotA |
| L S_circular_insula_sup | GM Secondary somatosensory cortex / Parietal operculum OP3 L | SomMotB |
| R G_and_S_subcentral | GM Primary somatosensory cortex BA3b L | SomMotB |
| R S_central | GM Primary motor cortex BA4p L | SomMotB |
| R S_central | GM Primary motor cortex BA4p R | SomMotA |
| R G_and_S_subcentral | GM Primary auditory cortex TE1.2 R | SomMotB |
| R G_Ins_lg_and_S_cent_ins | GM Insula Ig2 R | SomMotB |
| L G_front_middle | GM Broca's area BA45 L | ContA |
| L S_temporal_inf | WM Optic radiation L | DorsAttnA |
| L Lat_Fis-ant-Horizont | GM Broca's area BA44L | SalVentAttnB |
| L G_Ins_lg_and_S_cent_ins | WM Uncinate fascicle L | SalVentAttnA |
| R S_circular_insula_sup | GM Broca's area BA44 R | SalVentAttnA |
| L S_circular_insula_inf | GM Insula Id1 L | SalVentAttnA |
| R G_insular_short | WM Inferior occipito-frontal fascicle R | SalVentAttnA |
| L G_parietal_sup | GM Superior parietal lobule 7PC L | ContA |
| R S_interm_prim-Jensen | GM Anterior intra-parietal sulcus hlP1 R | ContA |
| L S_intrapariet_and_P_trans | GM Anterior intra-parietal sulcus hlP1L | ContA |
| L S_intrapariet_and_P_trans | GM Anterior intra-parietal sulcus hlP3 L | DorsAttnA |
| R S_intrapariet_and_P_trans | GM Anterior intra-parietal sulcus hlP3 R | DorsAttnA |
| R S_intrapariet_and_P_trans | GM Anterior intra-parietal sulcus hlP3 R | DorsAttnA |
| R G_insular_short | GM Broca's area B A44 R | SalVentAttnB |
| L G_temporal_middle |  | DefaultB |
| R G_temporal_middle |  | DefaultA |
| R G_temporal_middle |  | TempPar |
| R G_temporal_middle | GM Visual cortex V5 R | DorsAttnB |
| L G_temp_sup-Plan_tempo | GM Inferior parietal lobule PFcm L | TempPar |
| R Lat_Fis-post | GM Secondary somatosensory cortex / Parietal operculum OP2 R | SomMotB |
| R G_pariet_inf-Supramar | GM Secondary somatosensory cortex / Parietal operculum OP1 R | SalVentAttnA |
| L G_front_inf-Triangul | GM Broca's area BA45 L | DefaultB |
| L S_precentral-inf-part | GM Broca's area BA44 L | ContA |
| R G_front_inf-Orbital | GM Broca's area BA45 L | DefaultB |
| L Lat_Fis-post | GM Primary auditory cortex TE1.1 L | SomMotB |
| R G_temp_sup-G_T_transv | GM Primary auditory cortex TE1.0 R | SomMotB |
| L G_postcentral | GM Primary somatosensory cortex BA1 L | SomMotA |
| L S_postcentral | GM Inferior parietal lobule PFt L | DorsAttnB |
| L G_postcentral | GM Primary somatosensory cortex BA2 L | DorsAttnB |
| R S_postcentral | GM Primary somatosensory cortex BA2 R | DorsAttnB |
| L S_central | GM Primary somatosensory cortex BA3b L | SomMotB |
| L G_and_S_paracentral | GM Primary motor cortex BA4a R | SomMotA |
| L S_precentral-inf-part | GM Broca's area BA44L | ContA |
| R S_precentral-inf-part | GM Broca's area BA44R | ContA |
| R S_precentral-sup-part | GM Premotor cortex BA6 R | SomMotA |
| R S_precentral-inf-part | GM Premotor cortex BA6 R | ContA |
| L S_precentral-sup-part | GM Premotor cortex BA6 L | DorsAttnB |
| L S_precentral-sup-part | GM Premotor cortex BA6 L | DorsAttnB |
| L G_parietal_sup | GM Superior parietal lobule 5L L | DorsAttnB |
| R G_parietal_sup | GM Superior parietal lobule 7A R | DorsAttnA |
| R G_parietal_sup | GM Superior parietal lobule 7P R | DorsAttnA |
| L G_precentral | GM Primary somatosensory cortex BA2 L | SomMotA |
| L G_temp_sup-Plan_polar | GM Amygdala superficial group R | Limbic A |
| L S_temporal_transverse | GM Primary auditory cortex TE1.2 L | SomMotB |
| L G_temp _sup-Plan polar | GM Primary auditory cortex TE1.2 L | SomMotB |
| L S_temporal_sup |  | TempPar |
| R Lat_Fis-post | GM Insula lg1 R | TempPar |
| R G_temp_sup-Lateral | GM Insula ld1 R | TempPar |
| L G_and_S_subcentral | GM Inferior parietal lobule PFop L | SalVentAttnA |
| R S_postcentral | GM Inferior parietal lobule PFt R | DorsAttnB |
| L G_pariet_inf-Supramar | GM Inferior parietal lobule PF L | SalVentAttnB |
| S G_pariet_inf-Supramar | GM Inferior parietal lobule PF R | SalVentAttnB |
| L G_pariet_inf-Supramar | GM Inferior parietal lobule PF L | SalVentAttnB |

***Table S3. Regions related to MVPA analysis of different levels of the loudness feature.***

| Brain area | Hemisphere | Cluster size (mm^3^) | Z value | MNI coordinates | | |
| --- | --- | --- | --- | --- | --- | --- |
|  |  |  |  | x | y | z |
| Transverse temporal gyrus | L | 432 | 2.21 | -46.5 | -20.5 | 9.5 |
| Broca's area | L | 280 | 2.29 | -50.5 | 21.5 | 7.5 |
| Opercular cortex | L | 248 | 2.25 | -46.5 | -20.5 | 19.5 |
| Anterior supramarginal gyrus | L | 224 | 2.11 | -70.5 | -30.5 | 25.5 |
| Planum temporale (Sylvian fissure) | L | 168 | 2.20 | -44.5 | -38.5 | 11.5 |
| Anterior superior temporal sulcus | R | 128 | 2.17 | 43.5 | -38.5 | -0.5 |
| Planum temporale | R | 120 | 2.11 | 49.5 | -28.5 | 9.5 |
| Precentral gyrus | R | 96 | 2.10 | 59.5 | -6.5 | 45.5 |
| Posterior supramarginal gyrus | L | 96 | 2.23 | -50.5 | -46.5 | 13.5 |
| Central operculum | L | 96 | 2.17 | -54.5 | -18.5 | 13.5 |
| Posterior superior temporal sulcus | L | 96 | 2.16 | -52.5 | -10.5 | -20.5 |

***Table S4. Regions related to MVPA analysis of different levels of the tone feature.***

| Brain area | Hemisphere | Cluster size (mm³) | Z value | MNI coordinates | | |
| --- | --- | --- | --- | --- | --- | --- |
|  |  |  |  | x | y | z |
| Postcentral gyrus | R | 256 | 2.43 | 35.5 | -16.5 | 51.5 |
| Superior parietal lobule | L | 232 | 2.36 | -18.5 | -66.5 | 63.5 |
| Anterior supramarginal gyrus | R | 176 | 2.28 | 55.5 | -32.5 | 35.5 |
| Precentral gyrus | R | 176 | 2.43 | 29.5 | -16.5 | 57.5 |
| Planum temporale | L | 176 | 2.29 | -58.5 | -30.5 | 17.5 |
| Orbitofrontal cortex | R | 152 | 2.36 | 37.5 | 23.5 | -20.5 |
| Posterior angular gyrus | R | 128 | 2.14 | 35.5 | -70.5 | 51.5 |
| Supplementary motor area | L | 120 | 2.11 | -14.5 | -8.5 | 73.5 |
